# Single-cell temporal dynamics reveals the relative contributions of transcription and degradation to cell-type specific gene expression in zebrafish embryos

**DOI:** 10.1101/2023.04.20.537620

**Authors:** Lior Fishman, Gal Nechooshtan, Florian Erhard, Aviv Regev, Jeffrey A. Farrell, Michal Rabani

## Abstract

During embryonic development, pluripotent cells assume specialized identities by adopting particular gene expression profiles. However, systematically dissecting the underlying regulation of mRNA transcription and degradation remains a challenge, especially within whole embryos with diverse cellular identities. Here, we collect temporal cellular transcriptomes of zebrafish embryos, and decompose them into their newly-transcribed (zygotic) and pre-existing (maternal) mRNA components by combining single-cell RNA-Seq and metabolic labeling. We introduce kinetic models capable of quantifying regulatory rates of mRNA transcription and degradation within individual cell types during their specification. These reveal different regulatory rates between thousands of genes, and sometimes between cell types, that shape spatio-temporal expression patterns. Transcription drives most cell-type restricted gene expression. However, selective retention of maternal transcripts helps to define the gene expression profiles of germ cells and enveloping layer cells, two of the earliest specified cell-types. Coordination between transcription and degradation restricts expression of maternal-zygotic genes to specific cell types or times, and allows the emergence of spatio-temporal patterns when overall mRNA levels are held relatively constant. Sequence-based analysis links differences in degradation to specific sequence motifs. Our study reveals mRNA transcription and degradation events that control embryonic gene expression, and provides a quantitative approach to study mRNA regulation during a dynamic spatio-temporal response.

## Introduction

During development, transcript levels are tightly regulated by the combined action of mRNA transcription and degradation. By changing one or more of these processes, cells produce complex expression patterns that allow undifferentiated embryonic cells to establish distinct cell identities within the embryo. While the role of mRNA transcription in these events is very well established ^1, 2^, growing evidence highlights the importance of its precise interplay with mRNA degradation in shaping developmental gene expression patterns. For example, destruction of Nodal agonist and antagonist mRNAs triggered by the zebrafish microRNA miR-430 generates the precise expression levels needed for correct patterning of embryos ^3^. In addition, the localized stabilization of maternally inherited mRNAs within zebrafish primordial germ cells initiates a unique gene expression program ^4, 5^, that is only later supplemented by new zygotic transcription ^6^. However, despite its critical role in determining mRNA levels, the coordinated regulation of mRNA transcription and degradation remains less studied. It is still unclear to which extent does mRNA stability contributes to shaping developmental expression patters, and how does its coordination with mRNA transcription affects it. It is also unknown if and how regulation changes between genes and cell-types of a developing embryo undergoing cell-type specification.

Technical and computational challenges have limited the availability of genome-wide data on mRNA transcription and degradation during development. Conventional RNA-Seq ^7–9^, and more recently also single-cell RNA-Seq (scRNA-Seq) ^10–12^ measure overall mRNA levels, but provide only a partial view of mRNA dynamics. These approaches cannot distinguish between simultaneous transcription and degradation of transcripts in embryos. To address this challenge, RNA-Seq was combined with strategies for transcriptional arrest ^13^, quantification of intron-containing pre-mRNAs ^13, 14^, SNP detection ^15^ or RNA metabolic labeling ^16–18^. These strategies have generally used bulk sequencing, obscuring differences between individual cell types and diverse cellular identities within embryos. Recently, metabolic labeling of RNA was also combined with scRNA-Seq within cell culture contexts ^19–24^ and embryos ^25^, revealing cellular states and transitions at unprecedented resolution. However, such snapshot experiments are restricted in their ability to dynamically monitor changes in expression level and infer regulatory rates. Application of kinetic modeling tools has greatly enhanced single-cell studies ^19, 24^, and allowed to also monitor and quantify the underlying mRNA production and degradation rates in vitro. However, it is still unclear if such tools could be used to study whole embryos with diverse cellular identities and quantitatively separate the contribution of mRNA transcription and degradation in shaping spatio-temporal embryonic gene expression patterns.

The maternal-to-zygotic transition provides a compelling model to dissect the contributions of mRNA transcription and degradation to cell specification. At the onset of development, embryos are transcriptionally silent and rely on maternally-provided mRNAs and proteins that were deposited in the egg ^26^. In zebrafish embryos, this maternally controlled period lasts for approximately 3.3 hours, and involves many rounds of cell division. Once the maternal-to-zygotic transition begins in earnest, embryos undergo a massive degradation of maternally inherited mRNAs and begin to produce new zygotic transcripts ^26–28^. As the maternal-to-zygotic transition unfolds, cell-type specific changes in mRNA levels program pluripotent embryonic cells to assume more specialized identities ^2^. While transcriptional regulation is critical for these transitions ^2^, it has remained unclear to what extent regulation of mRNA degradation plays a role in these cell-type specification events. For instance, mRNAs for several lineage specific developmental regulators are also maternally deposited (e.g. *tbx16, cdx1b*, and others), and thus are initially ubiquitous. To direct downstream patterning, they must achieve a proper cell-type specific expression, which requires the destruction of non-specific maternal messages. However, how the interplay of mRNA transcription and degradation directs cell-type specific gene expression patterns within an embryo is still not resolved.

Here, we systematically quantify for the first time RNA transcription and degradation rates at cell-type resolution within a developing organism. We combine scRNA-Seq with RNA metabolic labeling within zebrafish embryos, and use it to follow the dynamics of the early zebrafish embryonic transcriptome. We decompose gene expression within single cells into its newly-transcribed (zygotic) and pre-existing (maternal) mRNA components, and develop dynamic models that allow us to resolve the relative contributions of mRNA transcription and degradation in determining spatio-temporal embryonic gene expression programs.

## Results

### Distinguishing newly transcribed from pre-existing mRNA within single cells of live embryos

We combined scRNA-Seq with RNA metabolic labeling to monitor newly-transcribed mRNA accumulation within single cells during zebrafish development (**Figure 1A**, **Methods**). We injected zebrafish embryos at the one-cell stage with 4sU-triphosphate (4sUTP), which is selectively incorporated into newly-transcribed RNA molecules and distinguishes them from pre-existing copies (**Methods**), which would have been maternally loaded. To benchmark our approach, we also injected into all embryos *in vitro* transcribed GFP and mCherry mRNAs that do not include any labeled residues (**Methods**). As was previously reported ^17^, injections did not interfere with normal zebrafish development phenotypically or transcriptionally. Injected embryos developed normally and reached the expected developmental stages. Single-cell transcriptomes obtained from these embryos are comparable to published ^11^ scRNA-Seq datasets of untreated zebrafish embryos (**Figure S1A**).

**Figure 1:**
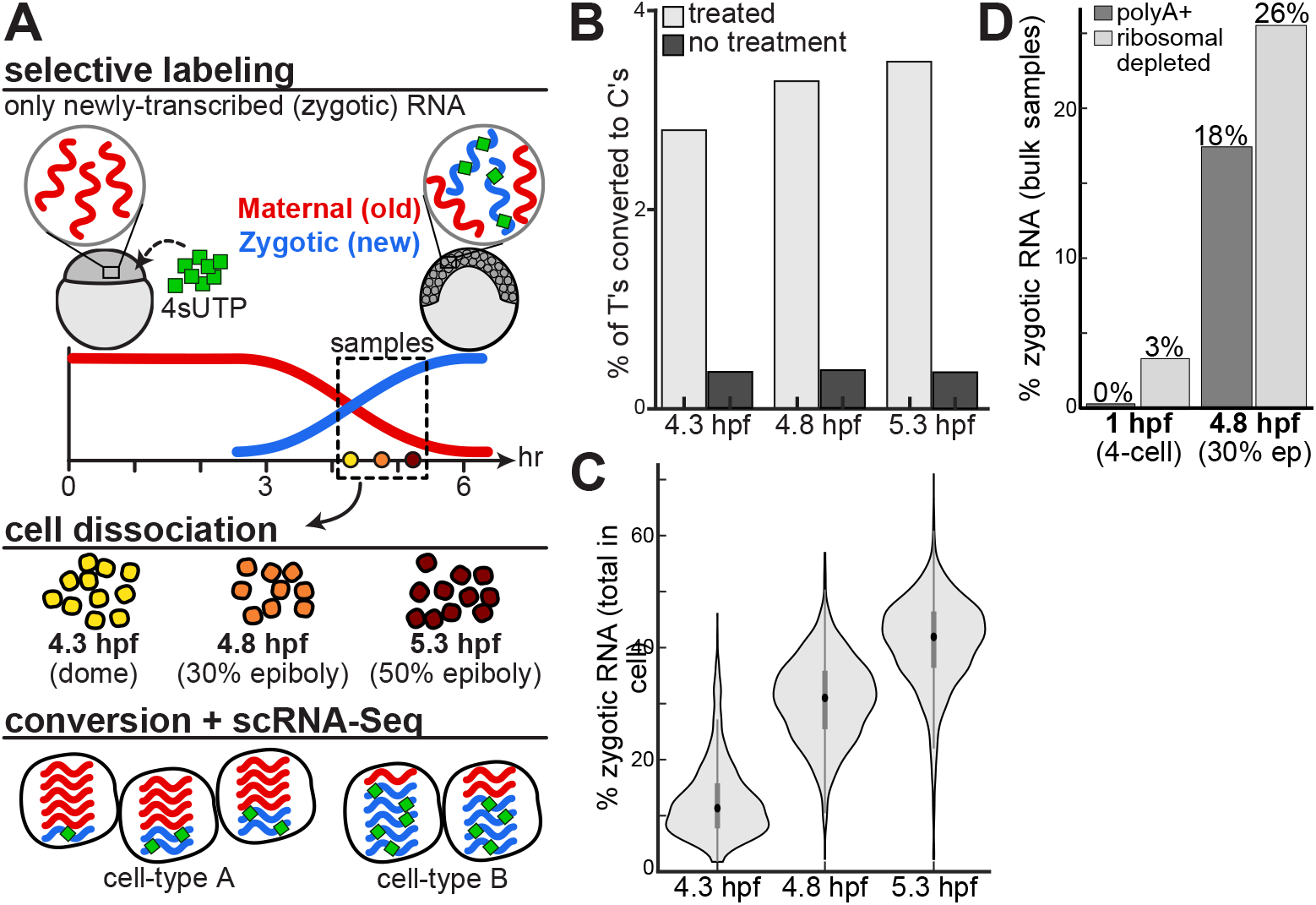
monitoring the embryonic maternal and zygotic transcriptomes at a single cell resolution. **(A)** An approach to combine scRNA-Seq with RNA metabolic labeling in zebrafish embryos. We injected zebrafish embryos at the one-cell stage with 4sU-triphosphate (4sUTP, green), which is selectively incorporated into newly-transcribed zygotic mRNA molecules (blue), while pre-existing maternally contributed molecules (red) remain unlabeled. We collected embryos at 3 developmental stages following the onset of zygotic transcription: dome (4.3 hpf; yellow), 30% epiboly (4.8 hpf; orange) and 50% epiboly (5.3 hpf; dark red). We dissociated embryos into single cells, and measured their transcriptomes by an adapted scRNA-Seq workflow that included a chemical conversion of labeled residues. Conversion induced T-to-C changes in downstream sequencing reads, enabling the separate quantification of newly-transcribed (zygotic) and pre-existing (maternal) mRNA within single-cell transcriptomes. **(B)** Fraction of T bases that were sequenced as C (y-axis) across all genes in the transcriptome, within each of 3 temporal samples (x-axis), when applying chemical conversion (light gray) or without such treatment (dark gray). **(C)** Distribution of GRAND-SLAM estimates of the overall percent of labeled RNA within each cell (y-axis) at 3 developmental stages (x-axis). The central dot is the median; the edges of the gray box are the 25th and 75th percentiles. **(D)** GRAND-SLAM estimates of percent of total labeled RNA per sample (y-axis) in bulk samples collected at two developmental stages using either polyA selection (dark gray) or ribosomal depletion (light gray).

To detect the incorporation of label within live cells, we adapted the Drop-Seq ^11, 29^ method by adding a chemical conversion of 4sU residues ^30^ after mRNA capture on beads (**Methods**). This conversion alters base pairing during reverse transcription and creates characteristic T-to-C changes in downstream sequencing reads, allowing quantification of newly-transcribed labeled RNAs. Post-treatment transcriptomes were comparable to controls that were not subjected to the additional conversion step for both native genes (**Figure S1B-C**) and injected controls (**Figure S1D**). These indicate that conversion did not interfere with the integrity of single-cell transcriptomes generated by Drop-Seq and did not increase barcode mixing between cells.

Using this approach, we collected 8,226 single-cell transcriptomes of live cells from embryos that were metabolically labeled starting at the one-cell stage (**Figure 1A**). We profiled embryos at 3 developmental stages following the onset of zygotic transcription (at 3.3 hpf): dome (4.3 hpf, 1,855 cells), 30% epiboly (4.8 hpf, 3,052 cells from 2 replicates) and 50% epiboly (5.3 hpf, 3,319 cells from 2 replicates). Indeed, the frequency of T-to-C conversion increased over developmental stages (**Figure 1B**), as expected when more newly-transcribed mRNAs have accumulated in cells.

During these developmental stages, embryos activate a massive degradation of maternally deposited mRNAs and simultaneously initiate new zygotic transcription. As labeling is applied immediately after fertilization, labeled nucleotides should only incorporate into newly-transcribed zygotic mRNAs, while pre-existing maternal transcripts that were produced prior to label injection should not show any significant labeling signal. To show the specificity of metabolic labeling to zygotic mRNAs, we analyzed a subset of known zygotic genes, and found high T-to-C conversion rates at all stages, as expected. On the other hand, conversion rates of both known maternal genes and injected controls remained low, at similar levels to untreated samples (**Figure S1E**).

We applied GRAND-SLAM analysis ^21^ to determine the fraction of newly-transcribed zygotic mRNA from T-to-C conversions for each gene in each cell. This approach deduces the fraction of labeled mRNA per gene from characteristically low 4sU incorporation rates (estimated 5.5% to 8.5% for zygotic genes in our samples, **Figure S1E**). It uses statistical inference to integrate the underlying position-specific incorporation rates, genetic polymorphism and other confounding effects, and improves accuracy of the estimated labeled fractions compared to the raw T-to-C conversion signals. Indeed, GRAND-SLAM analysis correctly estimated (**Figure S1F**) low labeling of pre-existing injected controls (labeled fraction < 0.8%, averaged across all cells) and known maternal genes (labeled fractions < 3.5%); as well as high labeling of newly-transcribed known zygotic genes (labeled fractions > 80%). The accuracy of estimated zygotic mRNA fractions was further improved by applying GRAND-SLAM to aggregated pseudo-bulk samples (**Figure S1G**), reaching nearly 100% for known zygotic genes (**Figure S1H**).

As expected, estimated labeled fractions within cells increased over developmental stages as more newly-transcribed zygotic mRNAs have accumulated in cells (**Figure 1C**). Labeled zygotic mRNAs accounted on average for only 13% of cellular mRNAs in early dome stage (4.3 hpf, **Figure 1C**), increasing to an average of 41% in late 50% epiboly stage (5.3 hpf). These results were also comparable to estimates by bulk RNA-Seq of metabolically labeled embryos. In bulk samples we detected only minimal levels of zygotic mRNAs in 1 hpf samples (4-cell, 0.3% in polyA+, 3.4% in Ribo-depleted, **Figure 1D**). However, the fraction of zygotic mRNAs increased substantially by 4.8 hpf (30% epiboly, 17.6% in polyA+, 26% in Ribo-depleted), and was similar to estimates in our single-cell analysis (31% on average at 4.8 hpf sample, **Figure 1C**).

Taken together, these results validate our method’s ability to accurately monitor newly-transcribed zygotic mRNAs and distinguish them from pre-existing maternal copies of genes in single cells of whole embryos. Our method provides a powerful and accurate approach that combines droplet-based scRNA-Seq and metabolic labeling to separately measure levels of maternally provided and zygotically transcribed mRNA within individual cells of a developing organism.

### Developmental pseudotime captures progression at the maternal-to-zygotic transition

To interpret single-cell transcriptomes, we used URD ^11^ to perform dimensionality reduction, UMAP projection, and clustering without considering whether mRNAs were maternal or zygotic. This analysis partitioned cells into 15 clusters (**Figure S2A**) reflecting both developmental stage and cell-type. Clusters were annotated based on their expression of known cell-type specific genes and visualized on a UMAP projection (**Figure 2A-C**). The UMAP represented well both the known specification events that have occurred at these stages of development (**Figure 2A**), and the temporal ordering of the 3 sampled timepoints (**Figure 2B**).

**Figure 2:**
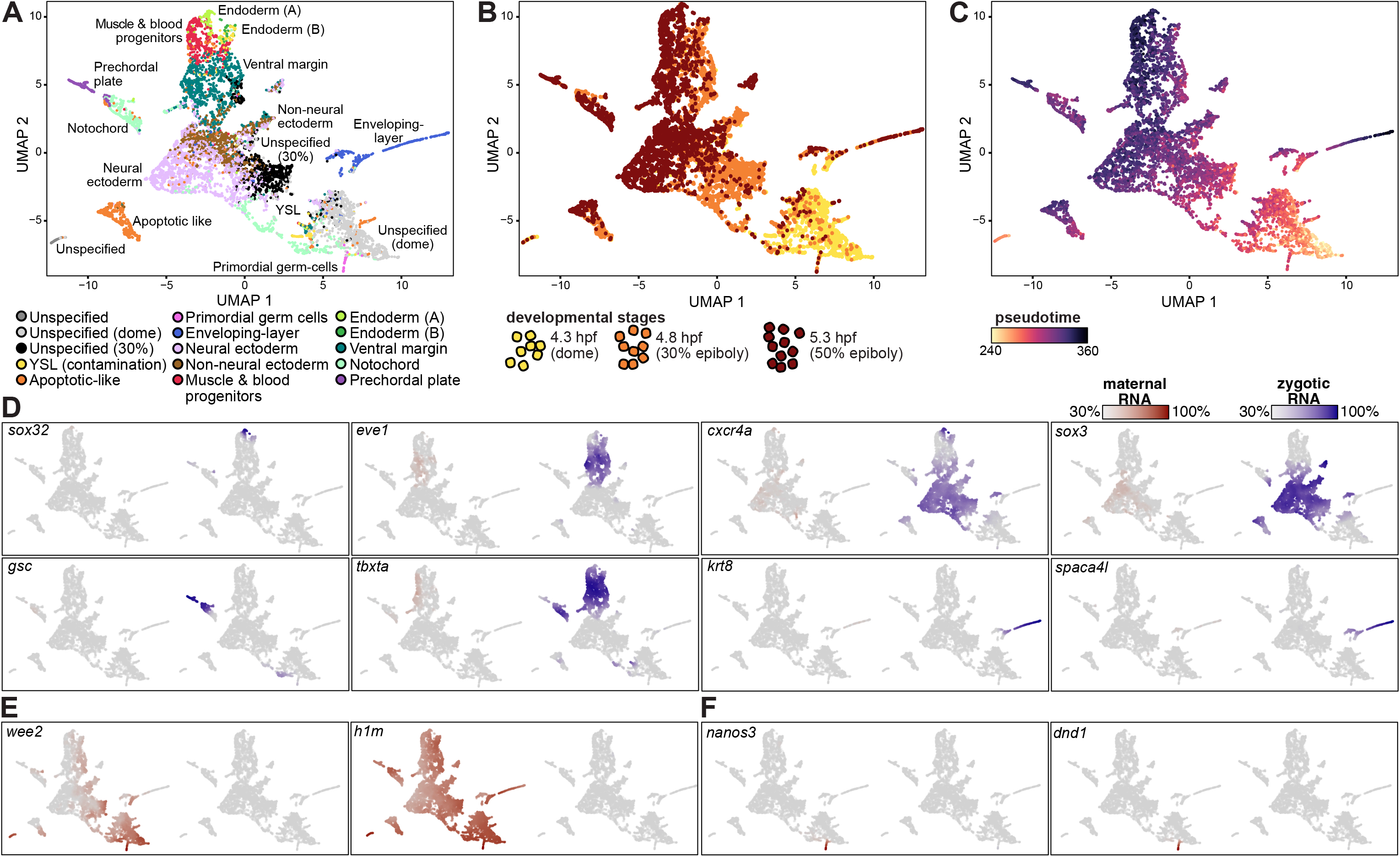
distinguishing maternal and zygotic expression across single cells of zebrafish embryos. **(A-C)** UMAP projection of 8,226 single cells from 5 embryonic samples (4.3 hpf, 1,855 cells; 4.8 hpf, 3,052 cells from 2 replicates; 5.3 hpf, 3,319 cells from 2 replicates). **(A)** Colored by 13 distinct cell-type clusters. 3 groups of unspecified cells do not express any specific cell-type markers, and represent cells which have still not differentiated. Two of these groups mostly include cells from one specific developmental stage, as indicated. Cells labeled as YSL are likely a contamination that originated from the yolk syncytial layer. **(B)** Colored by developmental stage: 4.3 hpf (dome, 1,855 cells, yellow); 4.8 hpf (30% epiboly, 3,052 cells, 2 replicates, orange); 5.3 hpf, (50% epiboly, 3,319 cells, 2 replicates, dark red). **(C)** Colored by pseudotime. Values range from 240 pseudo-min (earliest, yellow) to 360 pseudo-min (latest, purple). Pseudotime was calculated using the URD algorithm, based on the cells’ transcriptomic differences from the early unspecified cell populations. **(D-F)** Single cell expression of developmental genes across all 8,226 collected single cells. All cells are plotted, and each cell is colored by the normalized expression of a gene’s pre-existing maternal copies (red, left map) or newly-transcribed zygotic copies (blue, right map). The characteristically low label incorporation rates in combination with low per-cell number of reads by scRNA-Seq, limited the accuracy of estimated labeled mRNA fraction within single cells, which was often lower than expected within single cells and resulted in an unlabeled mRNA background. Therefore, color-scale of each gene is scaled by its maximal total expression, and its minimal 30% quantile of maternal and zygotic mRNA expression. Analyzed genes are indicated on plot. **(D)** Zygotically expressed genes, known to be specifically expressed within a certain lineage. **(E)** Maternally inherited genes. **(F)** Maternally inherited genes, known to be specifically expressed within germ-cells.

The levels of maternally deposited or zygotically transcribed mRNAs across different cells on the UMAP projection successfully recapitulated known developmental expression patterns. For example, zygotic transcription of cell-type specific zygotic regulators (**Figure 2D**) was identified in the mesoderm (*e.g., tbxta* and *gsc*), the ectoderm (*e.g., sox3* and *cxcr4a*), and in the enveloping layer (*e.g., krt8* and *spaca4l*). On the other hand, we recovered only pre-existing copies of maternally provided genes such as *h1m* and *wee2* (**Figure 2E**). Additionally, known maternal mRNAs that encode regulators of primordial germ cells (such as *nanos3* and *dnd1)* are appropriately restricted to this cell type (**Figure 2F**).

Since specification of cell-types in the blastula happens asynchronously, we used URD to compute a developmental pseudotime for each cell, which reflects each cell’s transcriptional difference from an early, unspecified cell population (**Figure 2C**). This calculation considered only total mRNA levels within cells. When considered separately, the decline of maternal and increase of zygotic mRNAs within cells were tightly associated with developmental pseudotime calculated on total mRNA. Differences in zygotic mRNA fractions between cells explained 70% of pseudotime differences (Pearson r^2^=0.7, **Figure S2B**). The early undifferentiated cells had few zygotically transcribed messages, whereas cells with later pseudotime had higher fractions of zygotically transcribed messages and corresponded to specified cell-types (**Figure S2C**). Cells in the enveloping layer had the highest zygotic mRNA fractions and pseudotimes (**Figure S2D**). These results demonstrate that developmental pseudotime accurately captures the accumulation of zygotic transcripts during development.

### Most embryonic genes are both maternally and zygotically contributed

Leveraging the ability of metabolic labeling to quantitatively distinguish the fraction of maternally contributed and zygotically transcribed copies of each gene, we analyzed the average fraction of newly transcribed zygotic mRNAs of each gene across single cells. We partitioned all 6,180 genes into 10 equally sized bins (quantiles) based on their average fraction of newly-transcribed zygotic mRNA in cells (**Figure 3A**).

**Figure 3:**
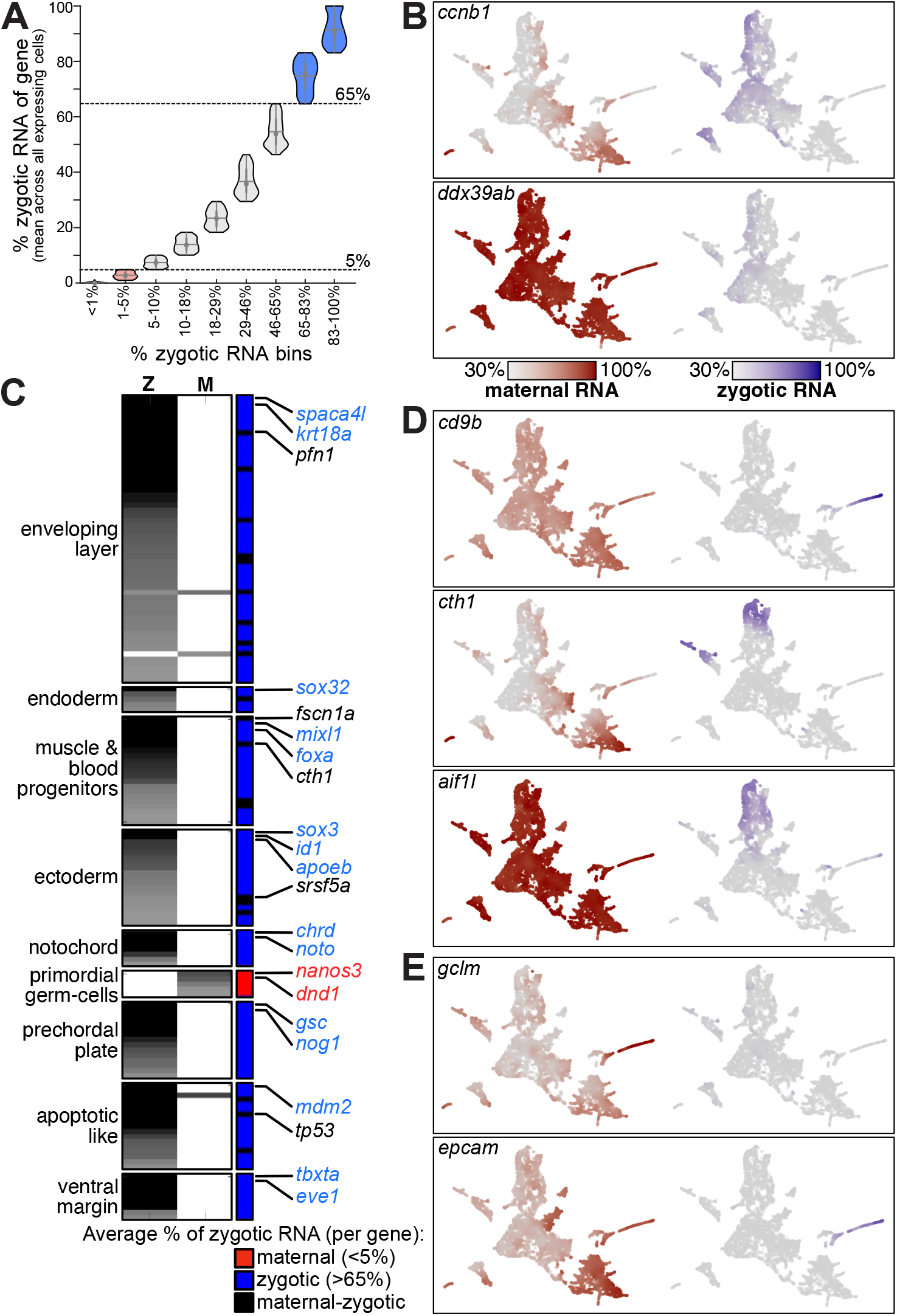
differences in zygotic mRNA accumulation between genes and cell-types. **(A)** Violin plots of the mean percent of zygotic mRNA per gene across all cells that express it (y-axis). A gene specific average fraction of zygotic RNA (across all cells) was calculated by averaging fractions across all cells which expressed it with 3 or more UMI counts. Genes were divided into 10 bins (x-axis), each with an equal number of genes. Since the fraction of zygotic mRNA per gene was equal (0%) for transcripts in the lowest two bins, they are plotted as a single set. The central dot is the median; the edges of the gray box are the 25th and 75th percentiles. We consider genes in the top two bins as zygotic (blue, zygotic mRNA fraction >65%) and genes in the bottom three bins as maternal (red, zygotic RNA fraction <5%). **(B)** Single cell expression of genes across all 8,226 collected single cells. All cells are plotted, and each cell is colored by the normalized expression of a gene’s pre-existing (maternal) copies (red, left map) or newly-transcribed (zygotic) copies (blue, right map). Color-scale of each gene is scaled by its maximal total expression and its minimal 30% quantile of maternal and zygotic mRNA expression. Analyzed genes are indicated on plot. **(C)** A set of 154 genes (rows) with either zygotically or maternally restricted expression within specific embryonic cell-types. Genes are ordered by cell-type, as indicated. Grayscale represents p-values for zygotic (left column) or maternal (middle column) RNA enrichment within cell-type (as indicated on left). Right column is colored by % zygotic RNA (red: <5%, maternal, 5 genes; blue: >65%, zygotic, 129 genes; black: maternal-zygotic, 20 genes). Selected gene ids are indicated on right (color code as indicated). Maternal-zygotic genes can be cell-type restricted in only one of the two components of their expression (e.g., *cth1* and *pfn1* are only cell-type restricted by their zygotic RNA, while their maternal mRNA is not restricted). **(D)** Single cell expression (as in B) of maternal-zygotic genes with cell-type restricted zygotic expression. **(E)** Single cell expression (as in B) of genes with cell-type restricted maternal expression in enveloping layer cells.

We considered genes within the lowest three quantiles of newly-transcribed mRNA (<5%) as maternal-only transcripts that are not zygotically transcribed during the first 5.3 hours of development. These included pluripotency and oocyte regulatory factors (*e.g., h1m, wee2*, **Figure 2E**), as well as known germ-cell regulators (*e.g., nanos3, dnd*, **Figure 2F**). This set is also specifically enriched for developmental processes involved in reproduction (p<4*10^-7^, hypergeometric Set Counts and Sizes (SCS) correction <5%), for endomembrane system (p<1*10^-28^) and for small molecule metabolism (p<2*10^-12^).

We considered genes in the top two quantiles of newly-transcribed mRNA (>65%) as zygotic-only transcripts that are not maternally inherited. These included many known zygotic developmental regulators (*e.g., gsc, cxcr4a*, **Figure 2D**) with documented functions within specific cell lineages. These zygotic-only transcripts were also enriched for transcription factors (p<2*10^-19^), patterning (p<9*10^-21^) and embryo development (p<4*10^-20^) annotations.

The estimated average fraction of newly-transcribed zygotic copies for the remaining 3,090 genes ranged between 5% and 65% (e.g., 47% in *ccnb1*; 8% in *ddx39ab*; **Figure 3B**), suggesting they are both maternally contributed and zygotically expressed during the first 5.3 hours post-fertilization. Genes in this group were enriched for functions related to RNA metabolism, such as splicing (p<1*10^-40^), ribosome biogenesis (p<5*10^-3^) and pol-II transcription (p<2*10^-3^), as well as other key cellular processes, such as translation (p<9*10^-48^), cell-cycle (p<4*10^-29^) and DNA replication (p<5*10^-17^). Components of these basic functions are both maternally contributed and expressed zygotically immediately following genome activation. Our data shows differences in both rates of maternal mRNA destruction and zygotic mRNA accumulation between transcripts in this group (**Figure 3B**). This suggests that both regulatory processes could serve to control expression of these genes during cell-type specification. However, in the absence of properly distinguishing maternal and zygotic transcripts as we do in this work, these dynamics would be obscured within total mRNA levels for most genes in zebrafish embryos, which have both maternal and zygotic contributions (**Figure 3A**).

### Zygotic transcription is the primary source of cell-type specific expression in embryos, and selective RNA stabilization is a minor source

During the maternal-to-zygotic transition, cell-type specific expression patterns emerge for many genes. Cell-type specific expression levels could be achieved in two ways at the maternal-to-zygotic transition: genes could be differentially transcribed in different cell types, or ubiquitous maternally deposited mRNA could be differentially degraded in different cell types. To investigate the prevalence of these different modes of regulation, we tested each gene for the enrichment of either its zygotic (newly-transcribed) or maternal (pre-existing) expression within different cell types (**Methods**). Overall, 436 of 6,180 genes (7%, **Table S1**) had either zygotically or maternally restricted expression within one or a few cell types (Kolmogorov-Smirnov Bonferroni <1%). Due to the high background in single-cell decomposition of maternal and zygotic mRNA, we conservatively limited further analysis to the 154 of those genes with the highest significance for cell type specificity (Kolmogorov-Smirnov p<10^-20^, **Figure 3C**).

Though most genes that are expressed in the embryo were both provided maternally and expressed zygotically, genes with cell-type restricted expression were overwhelmingly zygotic-only (129 genes, 84%, **Figure S3A**), including most well-known marker genes, such as *tbxta, noto* and *eve1* in the mesoderm, *sox17* and *her5* in the endoderm, and *krt5* and *krt8* in the enveloping layer. A much small number of maternal-only genes (5 genes, 3%, **Figure 3C**) are restricted to a specific cell type. All 5 of these maternal-only genes are restricted to the primordial germ cells and are well-established germ-cell markers^31^ (e.g., *dnd1, nanos3*, **Figure 2F**). We did not identify any additional zygotic markers within this cell type at those stages, consistent with prior findings that these cells are transcriptionally quiescent until later in development. Of these 5 markers, *dnd1, nanos3,* and *ddx4* have been shown to be selectively stabilized within zebrafish primordial germ cells^4, 5^, which is likely the case for the other 2 genes as well (*gra* and *ca15b*). This highlights selective stabilization of maternal mRNAs as the main mechanism for primordial germ cell specification during blastula stages.

Finally, we also identified a group of maternal-zygotic genes (20 genes, 13%, **Figure 3C**) with restricted expression within cell-types. In most cases, only the zygotic copies of these genes were restricted to specific cell types (e.g., *cth1* and *aif1l* in muscle and blood, or *cd9b* in enveloping layer, **Figure 3D**), while their maternal copies were not restricted. This indicates that their cell-type specific patterns are shaped largely by zygotic transcription rather than by cell-type specific stabilization of maternal mRNA. In these cases, the cell-type specific expression pattern of the zygotic copies is initially obscured by ubiquitously distributed maternal copies. Some genes exhibit quick elimination of maternal mRNA so that their cell-type specific expression pattern becomes clearly evident even during blastula stages (e.g., *cth1, cd9b*, **Figure 3D**, **Figure S3B**). Conversely, some genes exhibit slow degradation of maternal mRNA (e.g., *aif1l*, **Figure 3D**, **Figure S3C**), such that their cell-type specific transcription is obscured by remaining maternal copies of the same gene until later in development. Notably, two genes with both maternal and zygotic contributions exhibited cell-type specific enrichment of their maternally contributed copies – the enveloping layer-specific genes, *gclm* and *epcam* (**Figure 3E**). This suggests that these genes may achieve enveloping layer-specific expression during blastula stages through cell-type specific stabilization of maternal mRNAs.

Overall, this analysis supports the expectation that zygotic transcription is the main source of cell-type restricted expression during the specification of most cell types in zebrafish embryos. However, it also highlights the unique role of maternally inherited mechanisms in establishing the germ cell identity in zebrafish, and suggests that this mode of regulation also occurs within the enveloping layer for some genes. It is notable that the two cell types that exhibit a regulated stabilization of maternal mRNAs are the two earliest specification events during zebrafish embryogenesis ^11^. This aligns with the intriguing possibility that zygotic regulation is too slow to achieve patterning of these cell types sufficiently early in development.

### Kinetic models quantify regulatory parameters that govern gene expression changes in embryos

We aimed to obtain a more quantitative view of the contributions of regulation of maternal and zygotic transcripts to overall mRNA levels during cell-type specification, by modeling the dynamics of maternal and zygotic transcriptomes in our data (**Figure 4A**). To do so, we built kinetic models which leverage that our data (1) measures quantitative gene expression values at single-cell resolution, (2) clearly distinguishes maternal and zygotic transcripts from the same gene, and (3) contains multiple time points during early development. These kinetic models effectively allow parameterization of the rates of destruction of maternal mRNA and accumulation of zygotic mRNA and facilitate quantitative comparisons between different genes and different cell types. These rates can be used to reveal regulatory functions that govern gene expression changes in embryos and generate a quantitative view of coordinated transcriptional and posttranscriptional events.

**Figure 4:**
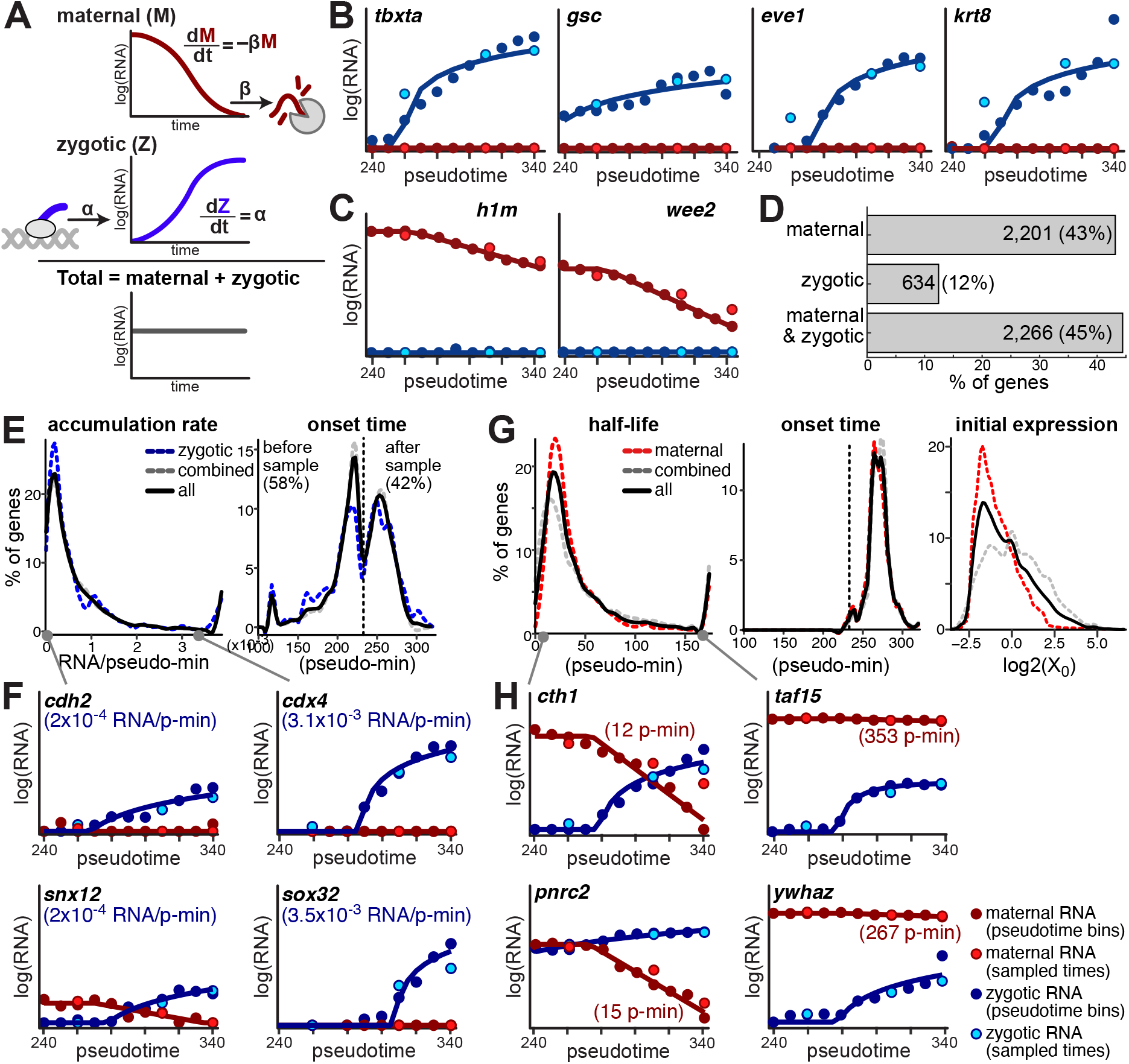
maternal and zygotic mRNA regulatory kinetic rates shape their temporal expression patterns. **(A)** A kinetic modeling approach to infer maternal and zygotic kinetic rates of each gene. Levels of maternal mRNA (M, red, top) are determined by an exponential decay of pre-existing copies with a constant degradation rate (β). Levels of zygotic transcripts (Z, blue, middle) are determined by a linear accumulation over time, with a constant accumulation rate (α). Total mRNA levels (bottom, gray) are the sum of both maternal and zygotic mRNA. We apply maximum-likelihood estimation to find the parameters (α, β) that best fit single cell observations. **(B-C)** Model fits (solid lines) to interpolated zygotic (blue dots) and maternal (red dots) expression levels (y-axis, log_2_ scale) across 11 pseudotime bins (x-axis) for key developmental genes. Gene name and predicted accumulation pseudo-rate are indicated on top. Light blue and light red dots represent estimated zygotic and maternal mRNA levels, respectively, for three sampled timepoints (4.3 hpf, 4.8 hpf and 5.3 hpf). **(B)** Zygotic regulators. **(C)** Maternally provided transcripts. **(D)** Histogram for number of genes (x-axis, fraction) that fit only a maternal model (top), only a zygotic model (middle) or both (bottom). Numbers and percentages are indicated. **(E)** Distribution of two parameters of the zygotic accumulation model across genes. Left: accumulation pseudo-rate (RNA/pseudo-min, x-axis) per gene (y-axis, fraction). Right: transcription onset time (pseudo-min, x-axis) per gene (y-axis, fraction). Time of earliest pseudotime bin in our data (240 min.) is indicated. Distributions are shown for all genes (black), zygotic only genes (blue) and combined maternal and zygotic genes (gray). **(F)** Model fits (as in B) for genes with very low (left) or very high (right) accumulation pseudo-rates. **(G)** Distribution of three parameters of the maternal decay model across genes. Left: degradation pseudo-rate (half-life, pseudo-min, x-axis) per gene (y-axis, fraction). Middle: degradation onset time (pseudo-min, x-axis) per gene (y-axis, fraction). Time of earliest pseudotime bin in our data (240 min.) is indicated. Right: initial expression level (log_2_, x-axis) per gene (y-axis, fraction). Distributions are shown for all genes (black), maternal only genes (red) and combined maternal and zygotic genes (gray). **(H)** Model fits (as in B) for genes with very short (left) or very long (right) half-lives.

First, we extrapolated high resolution expression dynamics from scRNA-Seq. Since specification of cell types in the blastula happens asynchronously, we used pseudotime (**Figure 2C**) to enable more detailed investigation of dynamics than could be achieved with 3 timepoints. We partitioned the cells into 11 pseudotime bins and estimated the zygotic (newly transcribed) and maternal (pre-existing) expression level of 5,101 genes within each temporal bin by pseudo-bulk analysis of all cells within a bin (**Methods**). Gene expression levels within bins were highly correlated to those calculated within 3 developmental stages (**Figure S4A**). Moreover, the high-resolution profiles recapitulated the expected expression dynamics of key genes. For example, we recovered upregulation of newly-transcribed mRNA of known zygotic regulators (**Figure 4B**), and downregulation of pre-existing mRNAs of known maternally provided genes (**Figure 4C**). This demonstrates the validity of our approach for interpolating the data between timepoints and calculating higher temporal resolution profiles.

We next developed a kinetic modeling approach to infer separate regulatory rates for maternal and zygotic copies of each gene. We estimate degradation pseudo-rates of maternal copies and accumulation pseudo-rates of zygotic copies per gene. Since our temporal information is based on a pseudotime analysis, the resulting pseudo-rates provide information on the relative production and degradation rates between genes in our dataset, rather than absolute measurements. We used a generative model (**Methods**, **Figure 4A**), in which levels of maternal mRNA are determined by an exponential decay of pre-existing copies (first order reaction), and those of zygotic transcripts are determined by a linear accumulation of new copies (zero order reaction), assuming minimal degradation. An alternative model which also includes a term for degradation of zygotic mRNAs could significantly improve the fit of only a small fraction of genes (261 genes, 9%, e.g., *cct6a* and *pum1*, **Figure S4B**), supporting a minimal effect of degradation of zygotic copies on overall mRNA levels within the timeframe of our experiment. We also assumed that, after onset, transcription and degradation rates are constant within the experimental timeframe (spanning less than 2 hours of development), but incorporated a gene-specific time of onset for each rate.

We applied maximum-likelihood estimation (**Methods**) to find the parameters of mRNA production and degradation profiles that best fit the dynamic observations and validated it using simulation studies (**Figure S5A-B**). Using a ‘goodness of fit’ test (**Methods**), the fitted kinetic models were retained for 97% of genes (4,923 genes at chi-square p>0.05, **Figure S5C**). As evidence that the determined rates are meaningful, the estimated pseudo-rates correlated to fold-changes measured in bulk SLAM-Seq samples (Pearson correlation > 0.43, **Figure S5D**). Additionally, the predictions from this kinetic model explained >96% (R-squared) of the variability in new RNA, old RNA and total RNA expression levels in our data, suggesting that these models with minimal parameters are sufficiently complex to capture the dynamics of gene expression in the early embryo. Altogether, these establish the ability of our kinetic models to infer kinetic rates of mRNA transcription and degradation from single-cell metabolic labeling data and successfully predict dynamic expression changes during a dynamic spatio-temporal response.

### Differences in regulatory rates between genes shape their temporal expression patterns

Our earlier analyses had revealed qualitatively that different maternal transcripts must be degraded at different rates (*e.g.* comparing *cth1* and *aif1l* turnover, **Figure 3D**). However, using parameters inferred by our kinetic models, we systematically compared all genes measured in this work to study the regulatory differences between them in their transcription, degradation and the coordination between them. Modeling results refined and extended the classification of genes. A subset of 2,201 genes (43%, **Figure 4D**) fitted only a maternal degradation model, suggesting these are predominantly maternally contributed with minimal evidence for their zygotic transcription within our data. Another 634 genes (12%, **Figure 4D**) fitted only the zygotic accumulation model, suggesting they are zygotically transcribed, with no evidence for any significant maternal contribution. Finally, 2,266 genes (45%, **Figure 4D**) fitted both models, suggesting these genes are both maternally provided and zygotically expressed during our measurement window. Overall, our models estimated degradation of maternal copies for 4,467 genes (88%) and accumulation of zygotic copies for 2,900 genes (57%).

Estimated zygotic accumulation pseudo-rates (2,900 genes, **Figure 4E**) varied by more than an order of magnitude between genes (median of 4.8*10^-4^ RNA/pseudo-min) and were highly correlated to expression levels. As expected from an accumulation model, faster production would result in accumulation to a higher level during a set time period. For example, genes such as *cdh2* and *snx12* accumulated more slowly (RNA/pseudo-min < 2*10^-4^, **Figure 4F**) while genes such as *cdx4* and *sox32* accumulated faster (RNA/pseudo-min > 3*10^-3^, **Figure 4F**). Our model also suggests that for a significant number of genes (1,608, 55%, **Figure 4E**) accumulation of zygotic copies started before our sampling window (onset < 240 pseudo-min). We did not observe any significant differences in accumulation rates between zygotic-only and maternal-zygotic genes. However, the 149 genes with cell-type restricted zygotic mRNA expression (**Figure 3C**), had higher than average accumulation rates (median of 1.7*10^-3^ RNA/pseudo-min).

Estimated maternal pseudo-half-lives of different genes (4,467 genes, **Figure 4G**) ranged within two orders of magnitude (median of 32 pseudo-min). For example, genes such as *pnrc2* and *cth1* decayed quickly (half-life < 15 pseudo-min, **Figure 4H**) while genes such as *taf15* and *ywhaz* decayed slowly (half-life > 250 pseudo-min, **Figure 4H**). On average, pseudo-half-lives of transcripts of maternal-only genes were slightly shorter than maternal transcripts of maternal-zygotic genes (61 pseudo-min. and 75 pseudo-min. respectively). Initial levels of maternal-only genes were also 2-fold lower on average than those of maternal-zygotic genes, while onset time of their decay was similar (263 pseudo-min on average).

These represent the first quantitative measurements of the degradation rate of maternal transcripts when their destruction is simultaneously obscured by zygotic replacement. Lack of notable global distinctions between maternal-only or zygotic-only genes and genes that are both maternally and zygotically expressed, suggests that similar pathways act to regulate transcription and degradation in all groups. Transcription and degradation rates can be 10-fold or more different between genes, highlighting their fine-tuned and precise regulation.

### Coordinated transcription and degradation of embryonic transcripts allow the emergence of spatially restricted patterns while total expression levels remain similar

For the 2,266 genes (45%) that are both maternally provided and zygotically expressed in our experiment, mRNA levels are affected both by degradation of pre-existing copies and accumulation of new ones. Interestingly, although mRNA levels of zygotic-only genes increased (positive fold-change) and those of maternal-only genes decreased (negative fold-change), mRNA levels of genes with both maternal and zygotic expression had much lower fold change values (**Figure 5A**). However, the distribution of their corresponding separate maternal and zygotic mRNAs mirrored that of maternal-only and zygotic-only transcripts (**Figure 5A**). This suggests that many transcripts are undergoing regulated replacement to maintain overall expression levels as new zygotic transcripts replace degraded maternal ones. Moreover, it emphasizes that total mRNA levels in this group obscure the dynamics of that replacement. Labeling and downstream deconvolution of maternal and zygotic transcripts is necessary to properly measure them.

**Figure 5:**
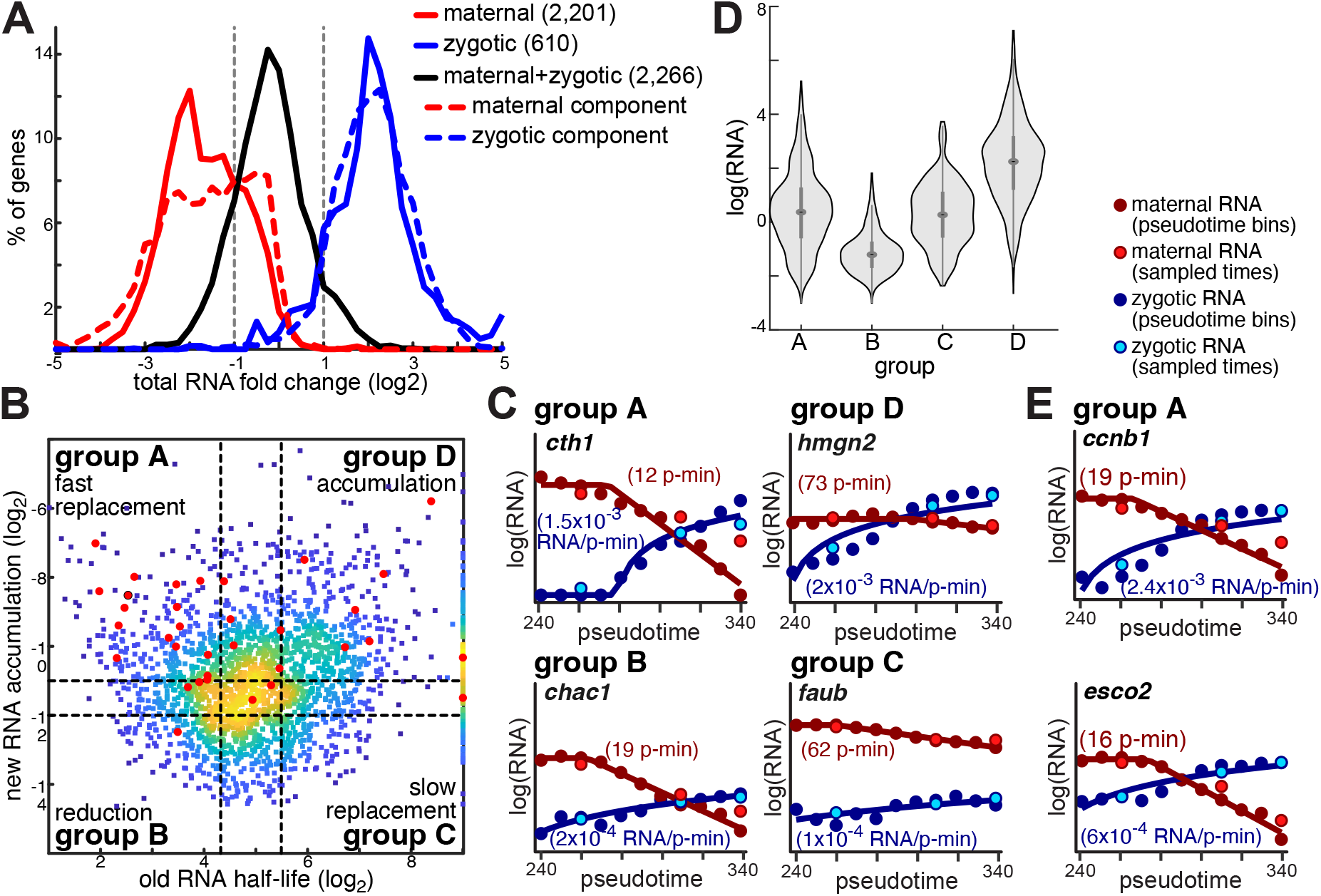
different replacement strategies between maternal and zygotic copies of embryonic genes underlie overall mRNA levels with fewer changes. **(A)** Distribution of total RNA fold-change (x-axis, log_2_) between average expression in 3 first pseudotime bins and 3 last pseudotime bins, of maternal-only (solid red), zygotic-only (solid blue) and maternal-zygotic (solid black) genes, as well as the separate maternal (dashed red) and zygotic (dashed blue) components of total maternal-zygotic RNA levels. Positive fold-change values represent an increase and negative values represent a decrease in mRNA levels over time. **(B)** Scatter plot of maternal mRNA half-lives (x-axis, log_2_) and zygotic mRNA accumulation (y-axis, log_2_) for maternal-zygotic genes. Colors represent density (yellow=high density; blue=low density). Red dots indicate genes that are included in the set of 154 genes with a cell-type specific expression. Dashed lines represent lower and upper bounds for defining four groups of genes within this set A-D (half-life bounds: < 20 pseudo-min or = 45 pseudo-min; log_2_ accumulation bounds: < -12 or = -11). **(C)** Model fits for one gene in each group (as indicated). Plots show model fits (solid lines) to interpolated zygotic (blue dots) and maternal (red dots) expression levels (y-axis, log_2_ scale) across 11 pseudotime bins (x-axis). Gene names, accumulation pseudo-rates and pseudo-half-lives are indicated. Light blue and light red dots represent estimated zygotic and maternal mRNA levels, respectively, for three sampled timepoints (4.3 hpf, 4.8 hpf and 5.3 hpf). **(D)** Distribution of total mRNA expression levels (y-axis, log_2_) for genes in each of the groups A-D (x-axis). The central dot is the median; the edges of the gray box are the 25th and 75th percentiles. **(E)** Model fits (as in C) for two genes with fast replacement in group A, whose expression is restricted to specific phases of the cell-cycle.

To analyze in depth the temporal dynamics of maternal-zygotic genes, we categorized genes into four groups (A-D, **Figure 5B**) based on the rate of destruction of their maternal copies and rate of accumulation of their zygotic copies (**Methods**). The 519 genes (23%) in group D combine slow destruction of maternal copies and fast accumulation of zygotic copies (e.g., *hmgn2*, **Figure 5C**), resulting in overall accumulation. This combination supports high expression levels for genes in this group, which are on average 4-fold higher than in other groups (**Figure 5D**). Genes in this group are enriched for RNA metabolic functions, such as RNA splicing (p<7*10^-35^) and translation (p<2*10^-52^), suggesting that embryos increase their capacity to process newly transcribed mRNAs as part of the process of initiating zygotic transcription. On the other hand, the 183 genes (8%) in group B, combine a fast destruction of maternal copies and slow accumulation of zygotic copies (*e.g., chac1*, **Figure 5C**), resulting in a decrease in total mRNA levels of genes in this group over time (1.7-fold decrease, on average), and overall lower expression levels (**Figure 5D**). This group is the only group enriched for components of the histone modification pathway (p<3*10^-7^).

Unlike expression levels of genes in groups B and D, which changed as a result of imbalanced destruction and replacement rates, expression levels of genes in group A and C remained relatively constant. The mean expression levels of genes in groups A and C are similar (**Figure 5D**), but their underlying replacement dynamics of maternal with zygotic transcripts differed markedly. Transcripts in group C (162, 7%) exhibited slow replacement, combining slow destruction of maternal transcripts with slow accumulation of their zygotic counterparts (e.g, *faub*, **Figure 5C**). Conversely, genes in group A (284, 13%) have fast replacement, combining fast degradation of maternal copies with fast accumulation of zygotic copies (e.g, *cth1*, **Figure 5C**).

While the slow replacement in group C represents an energetically favorable strategy, it is not clear why genes in group A are quickly replaced. We therefore tested for differences in the expression of genes that might suggest why. Indeed, group A is the only group enriched (16/34 genes, p<7*10^-7^, hypergeometric) with genes with cell-type restricted expression (**Figure 3C**). The combination of fast maternal degradation and fast zygotic accumulation in group A helps rapidly eliminate ubiquitous maternal expression and establish cell-type specific expression. Additionally, Group A is enriched for genes with tightly controlled temporal expression, such as cell-cycle genes (p<8*10^-5^), whose expression is restricted to specific phases of the cell-cycle (e.g., *ccnb1* at G2/M phase, *esco2* at S phase, **Figure 5E**). That we detect them as having fast replacement reflects that these genes are degraded and transcribed anew each cell cycle. Since cells in our data are not analyzed with regard to cell cycle progression, such genes will seem to be constantly expressed with high degradation and transcription rates.

Overall, this analysis reveals how coordination between transcription and degradation affects gene expression. For many genes, kinetics of replacement between maternal and zygotic mRNAs result in overall increases or decreases is total mRNA over time. But even when these processes are matched and genes maintain persistent expression levels, the underlying fast or slow rates have functional consequences for gene regulation. Slow rates conserve resources, but a rapid replacement helps to restrict expression of genes to a specific cell type or time.

### Differences in regulatory rates of embryonic genes between developmental trajectories identify lineage-specific mRNA degradation

Transcription rates often differ between cell types, contributing to cell-type specific gene expression patterns; a key question is whether maternal mRNAs exhibit different degradation rates between cell types that may also play such a role. We used URD to assign cells in our data to cell-type specific developmental trajectories. Then, we applied our kinetic modeling approach to infer maternal and zygotic regulatory rates along 5 separate cell-type-specific developmental trajectories, for which pseudotimes assigned to cells span a sufficiently large temporal interval (**Methods, Figure 6A, Figure S6A-B**). We used a ‘trajectory specific’ model that assumes that regulation can differ for a specific trajectory, and thus fits two separate sets of parameters for each gene: one to fit the ‘trajectory-specific’ profile, and another to fit all other cells not assigned to that trajectory. We compared this ‘trajectory specific’ model to a ‘uniform’ null model, which assumes similar rates across all cells. Genes that confidently (p < 0.05) reject the ‘uniform’ model in favor of the alternative ‘trajectory specific’ model represent cases where the dynamics of that mRNA are significantly different within one trajectory. In all other cases we retain the ‘uniform’ model. Given the lower numbers of cells within individual trajectories, we conservatively analyzed only a subset of genes with a significantly high expression within and outside each trajectory (**Methods, Figure S6C**).

**Figure 6:**
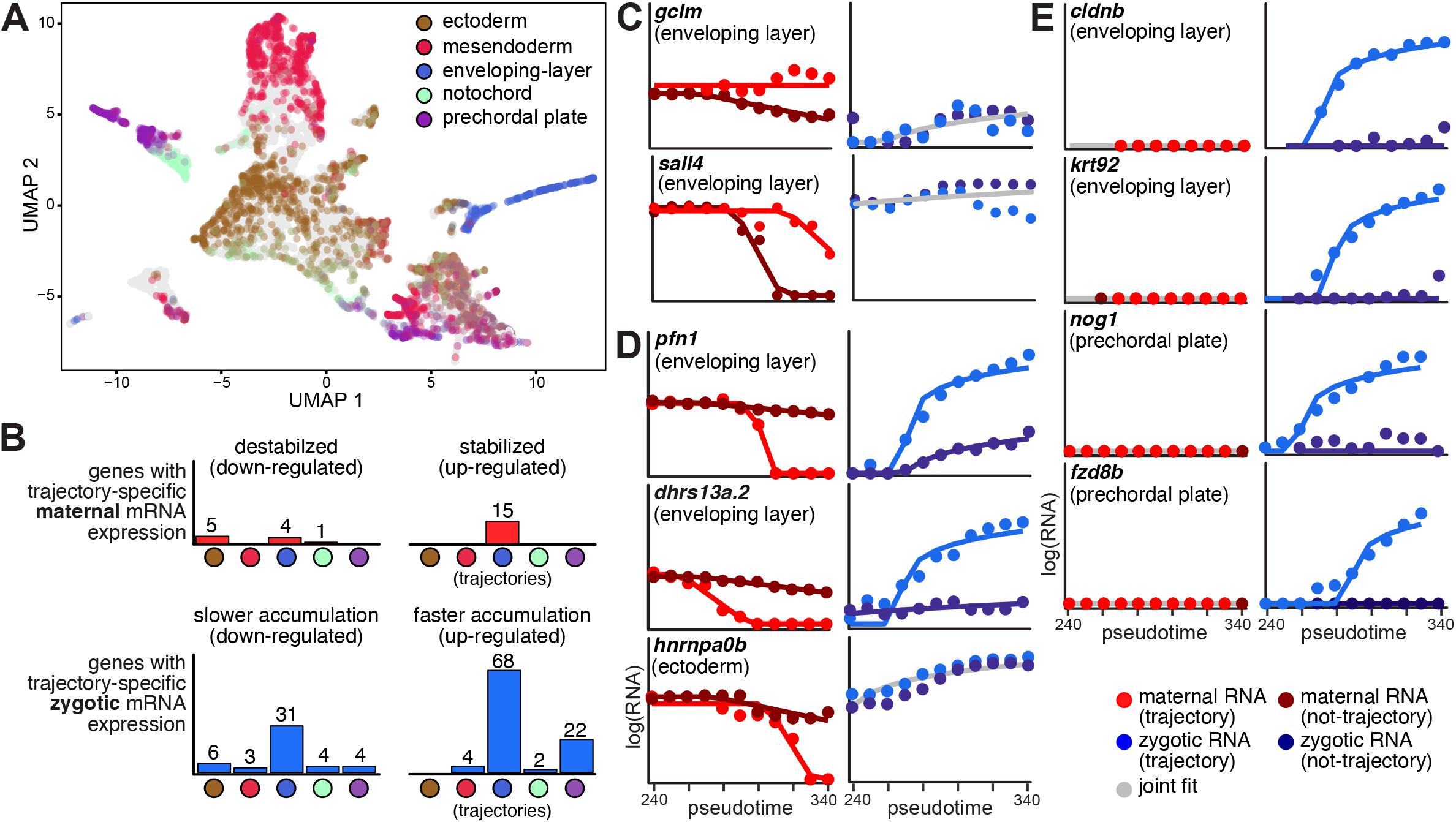
trajectory specific regulatory kinetic rates of embryonic genes. **(A)** A UMAP projection of 8,226 single cells with 5 cell lineages (colors as indicated). Each cell is colored by its lineage assignments. **(B)** Histograms (y-axis; number of genes) of trajectory-specific maternal (red, top) and zygotic (blue, bottom) significantly differentially regulated genes, per trajectory (x-axis, color-coded as in (A)). Each set of trajectory-specific genes is divided into up-regulated (right) and down-regulated (left) genes. Some genes are differentially regulated in more than one trajectory. **(C-E)** Trajectory-specific model fits (solid lines) to interpolated zygotic (right) trajectory (light blue dots) or non-trajectory (dark blue dots) and maternal (left) trajectory (light red dots) or non-trajectory (dark red dots) expression levels (y-axis, log_2_ scale) across 11 pseudotime bins (x-axis) for genes with trajectory specific regulation. Gene name and trajectory are indicated on top. Gray lines represent fits that match both trajectory and non-trajectory data, and therefore retain the null hypothesis of similar regulation within and outside a trajectory.

Most genes (2,196/2,345, 94%) retained the ‘uniform’ regulation hypothesis, suggesting that differences between cell lineages contribute minimally to shaping their mRNA dynamics. However, 149 genes had evidence for lineage-specific mRNA dynamics (**Figure 6B**, **Table S2**), indicating that lineage-specific regulation significantly contributes to shaping their expression levels, either through changes to maternal degradation or zygotic transcription.

When considering maternal regulation, we identify lineage-specific stabilization of 15 maternal genes. These were exclusively in the enveloping-layer (e.g. *gclm, sall4*, **Figure 6C**), supporting our previous observation for utilization of maternal stabilization in this cell-type, and identifying additional mRNAs that are differentially degraded in this cell type. Unfortunately, as the germ-cell lineage did not span a large enough temporal interval to be included in this analysis, we could not test our previous observations of maternal stabilization in germ cells in this analysis. Lineage-specific maternal destabilization was not investigated by our previous cell-type enrichment analysis (**Figure 3C**), and could represent an additional regulatory layer. We find evidence for maternal destabilization of 10 genes, including 4 in the enveloping-layer (e.g. *pfn1, dhrs13a.2*, **Figure 6D**) and 5 in the ectoderm (e.g. *hnrnpa0b*, **Figure 6D**).

When considering zygotic regulation, we find evidence for faster lineage specific accumulation of zygotic copies of 93 genes, including 59 out of 123 genes that we previously identified with cell-type specific expression in one of these lineages (**Figure 3C**). Faster accumulation is mostly evident in the enveloping-layer (68 genes, e.g. *cldnb, krt92*, **Figure 6E**), and prechordal plate (22 genes, e.g. *nog1, fzd8b*, **Figure 6E**). Another 43 genes showed evidence for slower accumulation of zygotic copies in specific lineages. These were mostly genes that are enriched in other lineages. Finally, 4 enveloping-layer genes show lineage-specific regulation of both their zygotic and maternal copies, including 3 genes with faster zygotic accumulation and faster maternal degradation in enveloping-layer (e.g., *pfn1, dhrs13a.2*, **Figure 6D**). These are examples of lineage-specific fast replacement between maternal and zygotic copies, which could help establish precise expression levels within specific cell types.

Together, these results reveal that most lineage-specific regulation of mRNA levels happens by zygotic transcription, as expected. However, in addition to the primordial germ cells (as previously discussed), it highlights differential regulation of maternal mRNAs in the enveloping-layer, by lineage-specific changes to their stability. This phenomenon has not been previously documented in enveloping layer cells, and suggests that lineage-specific alteration of maternal mRNA destruction may be shared by the earliest cell types specified in the zebrafish embryo ^11^.

### 3’UTR elements are associated with differences in maternal mRNA degradation in embryos

Taking advantage of the broad and quantitative measurements of mRNA degradation obtained by our models, we analyzed 3’UTR sequences of maternal transcripts, and associate sequences within them with differences in degradation. We systematically tested all 4- to 8-nt-long sequences (k-mers) within annotated 3’UTRs for their association with differences in pseudo-half-lives and onset times estimated by our kinetic models (**Methods**). We restricted our analysis both by significance (Kolmogorov-Smirnov FDR <1%) and effect size (normalized fold-difference).

Several expected ^32^ regulatory elements were enriched. For example, in transcripts with short half-life, we identified the miR-430 seed (GCACUU, p<6*10^-18^, **Figure 7A**), Pumilio binding sites (UGUAUAU, p<8*10^-3^, **Figure 7A**) and the AU-rich consensus (UAUUUA, p<2*10^-3^, **Figure 7A**). Within transcripts with delayed onset of degradation, indicative of early stabilization, we identified stabilizing poly-U signals (UUUUUUUU, p<2*10^-8^, **Figure 7B**).

**Figure 7:**
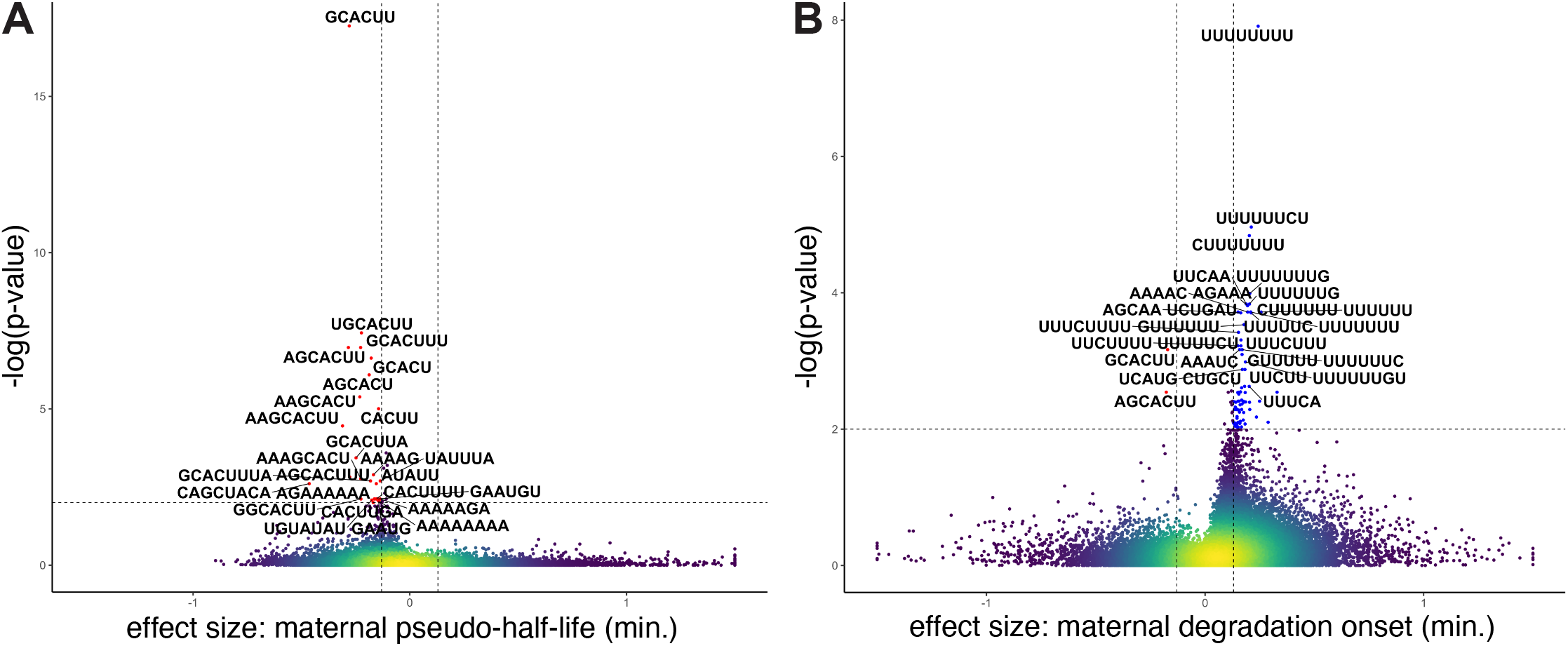
sequence enrichments associated with regulatory rates of mRNA degradation. Volcano plots for the enrichments of short sequences of length 5-8 bases (k-mers) based on difference between estimated parameters for genes that include this short sequence in their longest annotated 3’UTR sequence and those that do not. Plots show the significance (y-axis, -log_10_(p-value), Kolmogorov-Smirnov FDR < 1%) and the effect-size (x-axis, difference in the standard mean of each of the two distributions). Horizontal dashed line is 1% FDR, vertical dashed line is an absolute effect size of 0.13. **(A)** for the maternal pseudo-half-life parameter (log_2_) **(B)** for the maternal degradation onset time parameter. Colors represent density (yellow=high density; blue=low density). Dashed lines represent thresholds for significance. Top short sequences are noted on plots.

We also predict novel regulatory sequences within 3’UTRs. In particular, a delayed onset of degradation is associated with several A-rich sequences such as AGAAA (p<1.5*10^-4^, **Figure 7B**) and AAAAC (p<2*10^-4^, **Figure 7B**), as well as other sequences such as UUCAA (p<1.5*10^-4^, **Figure 7B**) and UCUGAU (p<2*10^-4^, **Figure 7B**). Interestingly, A-rich sequences are also associated in our analysis with a shorter half-life (AAAAG p<1.3*10^-3^, AGAAAAAA p<7.6*10^-3^, **Figure 7A**).

In addition to 3’UTR sequences, analysis of polyA tail lengths (as measured by ^33^) showed that longer half-lives are associated with longer tails as measured at 4hpf (p<6*10^-29^, Kolmogorov-Smirnov FDR <1%, **Figure S7A**).

We also analyzed 3’UTR sequences of maternal mRNA that were stabilized within germ cells. First, we reanalyzed our complete set of 436 cell-type specific genes, and identified 8 germ-cell enriched maternal transcripts (**Figure S7B-C**). Only one of these genes (*surf2*) showed any evidence of zygotic RNA transcription in our data. Interestingly, maternal copies of some of these genes (e.g., *dnd1, nanos3*, **Figure S7B**) are undetectable in single cell profiles of somatic cells, while other genes (e.g., *gra, ddx4*, **Figure S7C**) also show residual expression in somatic cells, as was also previously noted ^4, 34^. Analysis of published RNA-Seq data ^6^ of sorted germ-cell populations at dome stage (4.5 hpf) further confirmed that *nanos3* and *dnd1* are highly enriched in germ-cells (160 and 100-fold, respectively), while *ddx4* and *gra*, had a lower enrichment (45 and 29-fold, respectively). Therefore, we next searched for common sequences within the 3’UTRs of genes in each of these two groups. The miR-430 seed sequence (GCACUU) ^35^ occurred both in germ-cell specific (3/5 genes) as well as in those with residual somatic expression (1/3 genes). However, the Dnd1 binding sequence (UUUGAUUU) ^36^ occurred in 3’UTRs of all germ-cell specific genes (5/5 genes) but none of the genes with residual somatic expression. This suggests that protection by Dnd1 ^36^ is associated with maternal germ-cell markers that are completely depleted from somatic cells. However, other mechanisms to stabilize maternal transcripts in the germline, may work in a way that allows residual expression in somatic cells.

Finally, we looked for sequences that were over-represented in 3’UTRs of stabilized transcripts within the enveloping-layer. Interestingly, our analysis identified AU-rich elements (UAUUUAUU) as the most over-represented k-mers in this group (9/15 genes, p<1.5*10^-4^, Kolmogorov-Smirnov; due to the small number of genes, an FDR correction was not applied).

These results confirm the association of distinct regulatory elements with differences in regulatory pseudo-rates of embryonic genes, including both expected and novel signals. They distinguish two distinct residual expression patterns in somatic cells for maternal transcripts that are stabilized within the germline; but only one of which aligns with previously known sequence determinants of mRNA stability in those cells. These demonstrate our ability to utilize refined quantification by our kinetic modeling in order to generalize regulatory effects and identify sequence signals that encode them within 3’UTRs.

## Discussion

In this work we globally analyze mRNA transcription and degradation dynamics during cell-type specification in zebrafish embryos. We demonstrate the technological integration of metabolic labeling with scRNA-Seq in cells of a developing organism, sampled in a developmental timecourse. We develop kinetic models that integrate the analysis of timecourse data and generate a quantitative view of the relative contributions of mRNA transcription and degradation to gene regulation in the early zebrafish embryo. These reveal the regulatory functions and cell-type-specific variations that govern gene expression programs during a spatio-temporal dynamic response. Our portal (https://liorf.shinyapps.io/zebrafish_single_cell_regulation) provides the scientific community with ready access to our data and analysis results. We highlight three main regulatory principles.

Our models uncover fine-tuned and precise differences in regulation of mRNA transcription and degradation between genes that shape their spatio-temporal expression patterns. We generate for the first time precise and quantitative measurements of mRNA transcription and degradation rates of embryonic genes both globally and within individual cell types, which has been difficult to measure when their destruction is simultaneously obscured by zygotic transcription. As expected, most cell-type specific expression relies on the restricted zygotic transcription of genes. However, we find evidence for regulated stabilization of maternal mRNAs in two cell lineages: germ cells and enveloping layer cells. Both these lineages are specified early during zebrafish development ^11^, when transcription is not yet available as a tool to establish cell-type specificity. The yolk syncytial layer is another layer that forms similarly early in development and might have similar regulation of maternal RNAs. This tissue was not profiled in our study, but future single-nucleus RNA-Seq could allow study of this syncytial tissue, which is not divided into cells. Globally, regulation of maternal-only or zygotic-only genes is comparable to that of maternal-zygotic genes, suggesting that similar mechanisms regulate all different classes.

We show that coordination between transcription and degradation of maternal-zygotic genes maintains their overall mRNA levels, but that these parameters are tuned differently for each transcript. For example, slow exchange of maternal and zygotic copies of many housekeeping genes allows embryos to conserve and utilize pre-existing maternal mRNAs that are still needed later in development. On the other hand, fast exchange of maternal and zygotic copies helps to restrict zygotic expression to a specific cell type (e.g., *cth1* in muscle and blood, *cldn7b* in the enveloping layer) or time (e.g., *ccnb1* in G2/M phase of the cell cycle). Pre-existing maternal copies are targeted for fast removal, and allow quick establishment of cell-type or temporally restricted zygotic expression and eliminate background. A similar mode of regulation was also observed during fly development ^37^. It is unclear why a few cell-type specific genes implement slow exchange of maternal and zygotic copies, which obscures their cell-type specific expression until later in development (e.g., *aif1l* in muscle and blood). It was recently shown that mRNA destabilization in embryos can trigger compensatory changes in the transcription of homologues genes ^38^. It is intriguing to speculate that similar mechanisms could couple between transcription and degradation of maternally inherited mRNAs.

Finally, we associate known and novel elements within 3’UTRs of maternal mRNAs with differences in their degradation rate or timing as quantified by our models. For instance, miR-430 seeds, Pumilio binding sites, AU-rich elements, and A-rich sequences are all associated with shorter pseudo-half-lives; while poly-U signals, A-rich sequences, and some others (UUCAA, UCUGAU) are associated with delayed onset of degradation. These suggest a combined regulation of maternal mRNA stability by several pathways. We also analyze 3’UTR elements relative to cell type-specific changes in mRNA stability. We show that maternal germ cell markers with a Dnd1 binding site ^36^ are completely depleted from somatic cells (e.g., *dnd1, nanos3*). However, we also find markers with residual non-specific expression outside the germline (e.g., *gra, ddx4*). In those cases, other germ-cell RNA binding proteins such as Dazl-induced cytoplasmic polyadenylation ^39, 40^ could be involved. Interestingly, AU-rich elements were over-represented in 3’UTRs of stabilized transcripts within the enveloping-layer. Typically, AU-rich elements destabilize mRNAs, raising the possibility that this process is somehow attenuated within the enveloping-layer. Several RNA binding proteins exhibit enveloping layer-specific expression during blastula stages; among them, *rbm24a* has been linked to mRNA stabilization in zebrafish ^41^, suggesting it may play a role.

The technical framework of this work is reproducible and reliable by several tests, but could be further improved and expanded in certain directions. In particular, concentrations of 4sU that are tolerated by cells only replace a relatively low number of uridines by 4sU (at most 1 in 10 uridines in our samples, **Figure S1E**). Thus, statistical inference has more limited accuracy within single cells with fewer reads, and can underestimate labeled mRNA fractions in some cases. This problem is partially mitigated by performing pseudo-bulk analysis of scRNA-Seq data. Improved statistical modeling that aggregates similar cells within a sample could further reduce such biases. Interpolation of temporal information by pseudotime is also limiting, and allows inference of pseudo-rates rather than absolute time. Pseudotime units, which are estimated from transcriptomic changes, might also correspond to non-constant time units. However, as all genes will be similarly influenced by such biases, differences between genes will not be significantly affected. Optimizing the joint likelihood of rate parameters across all genes could further improve estimation accuracy. Finally, modeling within separate developmental trajectories is limited by temporal intervals. Measuring later developmental stages, when cell lineages have further differentiated, will allow more accurate quantification of differences in mRNA regulation. In addition, tools to control the timing of metabolic labeling within embryos will uncover the effect of mRNA stability of zygotic genes that drive later developmental events.

Our work systematically dissects quantitative mRNA transcription and degradation rates in vivo within developing embryos in an unprecedented cell-type and time resolution, and learns their regulatory principles that define and maintain gene expression programs during a dynamic spatio-temporal response.

## Acknowledgements

We thank Alex Schier for supporting early aspects of this project, for helpful discussions, and for providing access to sequencing facilities and zebrafish infrastructure. We thank Alex Schier, Eran Meshorer, Yotam Drier, Sagiv Shifman and Alon Zaslaver for critical reading of the manuscript. This research was supported by the Azrieli Faculty Fellowship and European Research Council Horizon 2020 grant 852451 (MR) and by the National Institutes of Health (K99HD091291 and ZIAHD008997 to JAF).

## Methods

### Zebrafish

All protocols and procedures involving zebrafish were approved by the Harvard University/Faculty of Arts and Sciences Standing Committee on the Use of Animals in Research and Teaching (IACUC; Protocol #25-08) and the Hebrew University Ethics Committee (IACUC; Protocol #NS-15859). Embryos were grown and staged according to standard procedures ^27^. Zebrafish embryos from wild-type AB/TL strains were used for all experiments.

### mRNA spike-ins cloning and transcription

Constructs encoding either a GFP or an RFP protein were PCR amplified and in-vitro transcribed from a T7 promoter with the HiScribe T7 Kit (NEB), using a mixture of ATP, CTP, GTP and UTP in transcription of GFP, or replacing UTP with 4sUTP (TriLink Biotechnologies) in transcription of RFP, to produce RFP mRNAs that were *in vitro* transcribed with 100% 4sU residues. Resulting mRNA levels were quantified by Qubit fluorometric quantification (Thermo Fisher) and mRNA length was validated by gel electrophoresis. For barcode mixing controls, mRNAs encoding mCherry with C-terminal fusions of either SV40 or nucleoplasmin nuclear localization signals were transcribed from pCS2-SV40nls-mCherry-SV40nls or pCS2-SV40nls-mCherry-NPLnls plasmids, using the mMessage Machine SP6 kit (Ambion/ThermoFisher).

### Fish microinjection and single-cell sample collection

Fertilized eggs were collected at 28°C, and kept in culture medium (5.03mM NaCl, 0.17mM KCl, 0.33mM CaCl_2_, 0.33mM MgSO_4_, 0.1% Methylene blue). One-cell staged wild-type zebrafish embryos were removed from their chorion and injected with 1nL of a solution containing 20mM 4sUTP (TriLink Biotechnologies), 30ng/uL GFP mRNA, 30ng/uL fully-labeled RFP mRNA and 5ng/uL of either mCherry-SV40nls or mCherry-NPLnls mRNA. For the first replicate, a total of 100 injected embryos were randomly collected at the Dome stage after visually ensuring that all embryos were at the same expected developmental stage. Embryos were placed in 2mL ice-cold deyolking buffer (55mM NaCl, 1.8mM KCl, 1.25mM NaHCO_3_), shaken in a thermal shaker (1100rpm) for 30 sec at 4°C and spun down (500g) for 1 min at 4°C. Cells were washed 3 times with 1mL ice-cold deyolking wash buffer (10mM Tris pH 8, 110mM NaCl, 3.5mM KCl, 2.7mM CaCl_2_)), shaken in a thermal shaker (1100rpm) for 30 sec at 4°C and spun down (500rpm) for 1 min at 4°C. Final pellet was resuspended in 500uL PBS. Cells were then passed through a 70µm cell sieve. Additionally, a total of 15-25 injected embryos were randomly collected per sample at the 30% and the 50% epiboly stages, and manually deyolked and dissociated as previously described ^11^. For the second replicate, a total of 70-75 injected embryos were randomly collected per sample at the 30% and the 50% epiboly stages, after visually ensuring that all embryos were at the same expected developmental stage. Embryos were then deyolked and dissociated similarly to dome stage embryos in the first replicate.

### Fish microinjection and bulk sample collection

AB/TL embryos were injected at the 1-cell stage with 1 nl of a solution containing 0.025% phenol red, 0.1 M KCl and 50 mM 4sU (Sigma). Embryos were grown in the dark and collected at the 4-cell and 30% epiboly stages (25 embryos each). Upon collection, embryos were disrupted in TRI Reagent (Sigma). RNA was isolated according to the manufacturer’s protocol, followed by ethanol precipitation. RNA was then treated with a TURBO DNA-free kit (Invitrogen) according to the manufacturer’s protocol, followed by ethanol precipitation. For RNA alkylation, RNA (2.9 µg) was first preincubated with 1 mM dithiothreitol (DTT) for 10 min at 55 °C and transferred to ice. Alkylation was conducted in sodium phosphate buffer (46.6 mM Na2HPO4, 3.4 mM NaH2PO4, pH 8), 50 % dimethyl sulfoxide (Sigma) and 10 mM iodoacetamide (Sigma; freshly prepared 100 mM stock dissolved in ethanol). The reaction was incubated 15 min at 50°C and quenched by adding DTT to 20 mM. RNA was cleaned up on an RNA Clean & Concentrator-25 column (Zymo) according to the manufacturer’s protocol. For poly(A)+ RNA sequencing, libraries were prepared using a KAPA Stranded mRNA-Seq Kit (Kapa Biosystems) according to the manufacturer’s protocol and sequenced on an Illumina NextSeq 500 platform with 75 nt single-end reads. Sequencing of ribo-depleted RNA was performed by Macrogen. Libraries were prepared with an Illumina TruSeq Stranded Total RNA with Ribo-Zero Human/Mouse/Rat kit, and sequenced on an Illumina NovaSeq 6000 platform with 100 nt paired-end reads. Data was deposited in GEO under accession GSE224113.

### Collection of Drop-seq transcriptomes

Drop-seq droplet encapsulation of cells was performed as previously described ^11, 29^. We added an RNA alkylation step ^30^ after mRNA capture on beads. Such chemically induced alkylation of 4sU residues alters base pairing during reverse transcription and creates characteristic T-to-C conversions in downstream sequencing reads, allowing quantification of labeled RNAs. Droplets were held on ice for up to 2 hours (to enable multiple sequential collections) before starting the RNA recovery, chemical treatment and reverse transcription. Libraries were built according to the Drop-seq protocol version 3.1 (12/28/2015, available http://www.dropseq.org/) with the following modifications. Iodoacetamide (IAA, Sigma-Aldrich) was freshly dissolved in ethanol to make a 100mM stock solution. Following breakage of droplets, beads were washed once with 300uL of 5x IAA buffer (250mM NaPO_4_ pH8), and incubated in IAA solution (50mM NaPO_4_ pH8, 10mM IAA, 20% DMSO, 6% Ficoll PM-400) for 15min at 32°C with rotation. DTT was added to a final concentration of 20mM to stop the reaction, and beads were washed twice with 6x SSC solution before continuing with reverse transcription according to standard protocol. An estimated 50–100 STAMPs were included in each 50uL PCR reaction, and 12–13 cycles of PCR amplification were performed. Libraries were sequenced using Illumina Nextseq v2.2 Mid or High Output 150bp chemistry. Data was deposited in GEO under accession GSE224918.

### Alignment and quality control of Drop-seq data

#### Processing of sequencing data

Alignment of sequencing reads and generation of digital expression matrices was performed essentially as previously described ^11^, using Drop-seq tools v1.12. Since the T-to-C conversions induced by alkylation of 4sU could affect mapping rates, we made modifications to ensure that T-to-C conversions did not reduce mapping rates by converting all bases in the genome and sequencing reads. Ensembl Zv10 Release 82 reference genome was used with transcripts assigned gene names by giving priority to names from ZFIN (as previously described ^11^, except with the reference modified to include our four spike-in sequences. Two modified versions of the genome were produced: one where all T residues were replaced with C and one where all A residues were replaced with G. Prior to alignment, original sequencing reads were preserved and then all A residues were replaced with G. These converted reads were aligned to both modified genomes using Bowtie2 as previously described, then filtered based on which strand the read was aligned to. These two outputs were combined and then processed through the remainder of the Drop-seq tools pipeline to produce processed BAM files and digital gene expression matrices. Converted reads were then replaced with the original reads and CIGAR strings were recalculated using Samtools in the final output BAM files. While a significant number of reads was mapped to 3 non-labeled controls (GFP and mCherry spike-ins sequences), we detected only very few reads that mapped to the 100% 4sU-labeled RFP injected control, which did not allow to infer conversion rates for this control. We indeed validated that the IAA treatment of this 100% labeled mRNA led to a significant degradation of this mRNA.

#### Identification of highly variable genes and batch correction

First, the digital gene expression matrix output by Drop-seq tools was imported into URD ^11^, filtered to retain only cells with expression of at least 500 genes, and the data was log-normalized. 924 highly variable genes were identified using findVariableGenes (*diffCV.cutoff = 0.2*). There was a noticeable batch separation between samples that received IAA treatment and control samples, as well as between samples processed using the two different dissociation methods described above. To correct for this, gene expression modules were calculated using non-negative matrix factorization as described (Farrell et al 2018 Science) with parameters (*k = 35, rand_state = None, alpha = 2, l1 = 0.5, max_iter = 10000, rep = 5, init = random*). 6 modules were identified that differed primarily between dissociation methods or treatment conditions (NMF 2, NMF 7, NMF 11, NMF 14, NMF 17, and NMF 22). Genes highly loaded into those 6 batch-associated modules (as determined by an elbow plot of gene loadings), as well as mitochondrial genes and spike-ins were removed from the highly variable genes, leaving 854 remaining highly variable genes.

#### Determination of cell clusters

Principal component analysis was computed using the highly variable genes (*calcPCA)*. Cell type clusters were calculated with Jaccard/Louvain graph clustering (*graphClustering, dim.use = “pca”, which.dims = c(1:7, 9:10, 12:16), method = “Louvain”, do.jaccard = T*). Principal components to include in the clustering were determined based on an elbow plot and by inspection of top loaded genes to identify PCs that represented known developmental processes in zebrafish blastula. PC8 was excluded because it was primarily associated with technical quality of libraries, and PC11 was excluded because it was primarily associated with cell cycle stage. Clusters were annotated based on expression of known marker genes (*nanos3, dnd1, krt8, gsc, noto, tbxta, eve1, chrd, sox3, isg15, sesn3, osr1)* and developmental stage. Endodermal progenitors were manually separated into their own cluster based on their expression of *sox32*.

#### UMAP projection

First, a diffusion map was computed using *destiny*, as incorporated into URD (*calcDM, knn = 50, sigma.use = ‘local’*). Then, the first 25 diffusion components were used as the basis to compute a UMAP projection using the *umap* function from the R package *umap*.

#### Pseudotime calculation

Pseudotime was calculated using URD (*floodPseudotime, root.cells = (see below), n = 100, minimum.cells.flooded = 2*). URD requires user specification of the youngest cells or ‘the root.’ 100 cells were chosen to use as the ‘root’ (i.e. the starting point for the pseudotime calculation) based on 3 criteria: (1) they were from the earliest stage in the data (dome), (2) they were in the dome stage cluster that did not exhibit signs of specification (cluster 10, “Unspecified Dome Stage”), and (3) they had the highest expression of an NMF module (NMF33) defined primarily by exclusively maternally loaded genes (e.g. *cldng, cldnd, ccna1,* and *cth1*).

#### Identification of developmental trajectories

For a subset of cell types, developmental trajectories were calculated using URD. URD requires that the terminal cells or ‘tips’ are defined by the user. Tips were identified from each terminal cell cluster by identifying the 12.5% of the 50% epiboly cells (minimum 25 cells) that were oldest in pseudotime. The transition matrix was biased (*pseudotimeDetermineLogistic, optimal.cells.forward = 25, max.cells.back = 50*) and random walks were simulated from each tip (*simulateRandomWalksFromTips, n.per.tip = 10000, root.visits = 1, max.steps = 1000*). For each cell, its maximum number of visitations by walks from a single tip were determined. Then, cells that were visited at least 50 times (i.e. by at least 0.5% of random walks) were associated with a trajectory if they were visited during walks from that cell type tip at least 50% as often as they were visited by walks from whichever cell type visited them the most strongly.

### Analysis of nucleotide conversion signals

GRAND-SLAM (GRAND3_3.0.0) ^21^ was used to calculate the fraction of labeled RNA (NTR) of each gene. For analysis of bulk RNA-Seq datasets, GRAND-SLAM was used with default parameters and a gedi (GRAND-SLAM_2.0.5f) genome reference was created using GRCz11 genome (GenBank:GCA_000002035.4) and Ensembl annotations release 102 ^42^. For single-cell analysis of scRNA-Seq datasets, GRAND-SLAM was used with default parameters, and gedi genome reference was created by manual additions of RFP, GFP and mCherry spike-in sequences to the GRCz11 genome. For pseudo-bulk analysis of scRNA-Seq datasets, GRAND-SLAM was used with the following non-default parameters: -*pseudobulkName clusters, -pseudobulkPurity 0, -pseudobulkFile <PB_FILE=*, using a different file for each of the different pseudo-bulk runs (cell-types, developmental stages or pseudo-time bins).

### Analysis of maternal and zygotic expression levels in single cells and UMAP visualization

Total normalized RNA expression per cell was calculated from the number of unique molecular identifiers (UMIs) using the *NormalizeData* function from Seurat version 4.1.0 ^43^ with default parameters. Briefly, normalization divides the total number of UMIs of each gene in the cell by the total UMI counts in the cell, multiplied by 10,000 and natural-log transformed using *log1p*. For a gene ! at cell *c* this is given by:

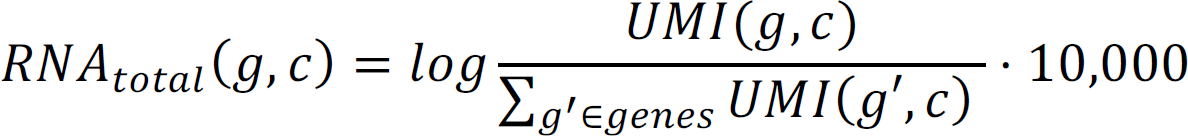

Fraction of labeled RNA (NTR) estimated by GRAND-SLAM per gene *g* in cell *c* was used to calculate maternal and zygotic RNA expression by:

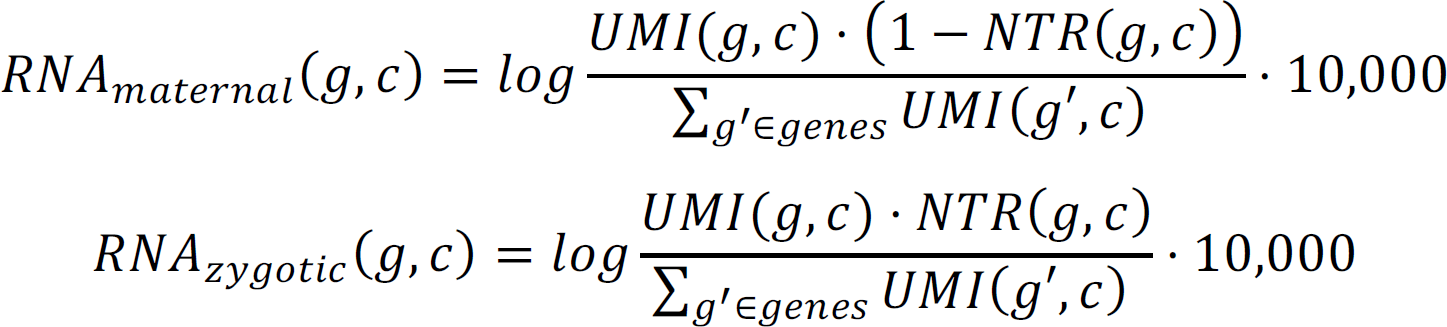

For plotting maternal and zygotic RNA expression on a UMAP projection, we first calculated a distance matrix based on the pairwise similarities between the cells’ embeddings. The similarities were calculated using a gaussian kernel function with a beta parameter, which adjusts the spread of the kernel. Each of the total-RNA, maternal-RNA or zygotic-RNA normalized single cell expression matrices were multiplied by the distance matrix and divided by the sum of expression in each row. A single UMAP color-scale was defined for each gene, and used for both maternal-RNA and zygotic-RNA plots. Color-scale maximum was defined by the maximal total-RNA density value. Color-scale minimum was defined by the 30th percentile of maternal-RNA and zygotic-RNA density values across all cells with a UMIs count of 3 or more for the gene.

### Analysis of maternal and zygotic expression levels in pseudo-bulk samples

Expression values (RNA) calculated by GRAND-SLAM were normalized as follows, for gene ! at pseudo-sample *s*:

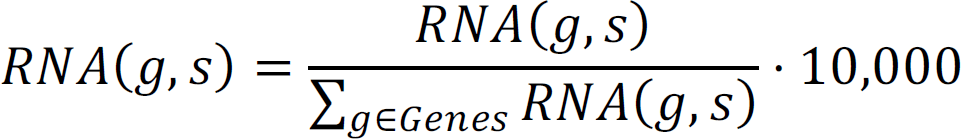

Fraction of labeled RNA (NTR) estimated by GRAND-SLAM per gene *g* in pseudo-bulk sample *s* was used to calculate maternal and zygotic RNA expression by:

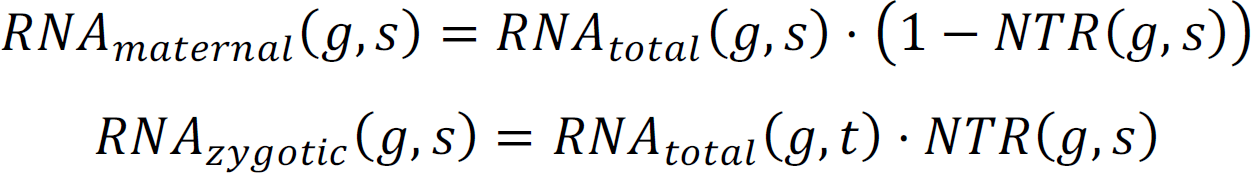

Final expression levels were log-transformed and floored to -4.

### Analysis of total expression levels in scRNA-Seq samples

For pairwise comparisons of single-cell samples, in each of the two samples compared, we calculated a pseudo-bulk expression level for each gene ! by:

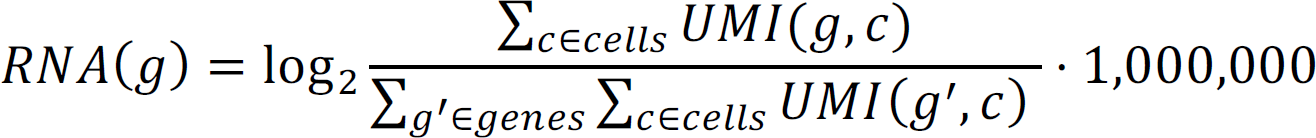

### Selecting genes with cell-type restricted maternal or zygotic mRNA expression

For each gene in our data, we tested the enrichment of either its zygotic (newly transcribed) or maternal (pre-existing) transcripts, by comparing the distribution of its maternal or zygotic UMI counts within cells assigned to a specific cell-type and cells not assigned to that type with a one-sided Kolmogorov-Smirnoff test, requiring higher counts within a cell-type. The maternal and zygotic UMI counts per cell were calculated by multiplying the total single-cell UMI count by the fraction of labeled RNA (NTR) as calculated by GRAND-SLAM for single cells. We restricted our analysis by pseudotime, and tested the enrichment of cells within a specific cell-type compared to other cells in our data of a similar pseudotime (difference of less than 20 pseudotime units). P-values were corrected for multiple hypothesis testing by a 1% Bonferroni correction. For each gene, we considered only the most significantly enriched cell-type for further analysis (after Bonferroni correction).

### Dividing cells into pseudotime bins

URD assigned pseudotime values (per-cell) were converted from arbitrary units (au) to pseudo-minutes post fertilization (pseudo-mpf) using a simple linear transformation, so that the first sample is assigned the value 240 pseudo-mpf and the last sample 360 pseudo-mpf. Cells with a pseudotime (au) smaller than 0.1 were treated as 0.1. Cells were divided into 10 pseudo-minute bins with 3 pseudo-minute overlap between adjacent bins. The last bin, which was significantly smaller than the rest, was joined with the preceding bin, creating 11 bins in total.

### Kinetic models of maternal and zygotic mRNA expression dynamics

We modeled maternal mRNA (1) and zygotic mRNA (Q) expression levels by two independent kinetic models.

#### Maternal mRNA

We compare three alternative models for temporal maternal mRNA kinetics. The simplest (null) model assumes no maternal expression of a gene, and has no parameters:

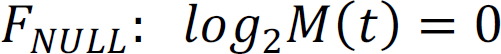

The simpler alternative model assumes very low degradation of maternal mRNA, and has only one parameter: initial expression level (W_X_),

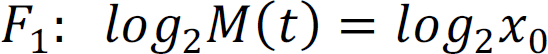

The second alternative model, describes dynamic changes in maternal mRNA levels using an exponential decay with 3 parameters: a constant decay rate (Z), initial expression level (W_X_), and degradation onset time (*d*):

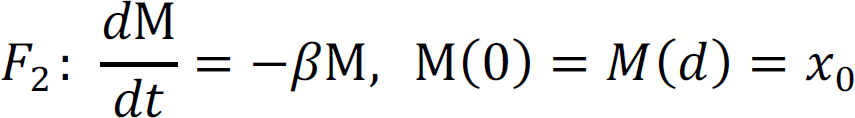

Solving this function analytically we get:

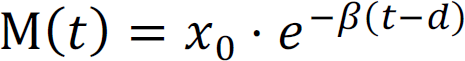

And we solve it looking for the log expression rate:

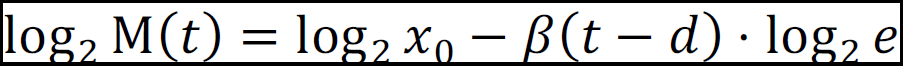

#### Zygotic mRNA

We compare three alternative models for temporal zygotic mRNA kinetics. The simplest (null) model assumes no zygotic expression of a gene, and has no parameters:

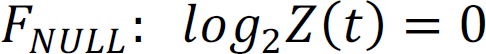

The simpler alternative model is a linear accumulation model with 2 parameters: constant transcription (a), and transcription onset time ([). This model assumes very low degradation of zygotic RNA during our sampling time.

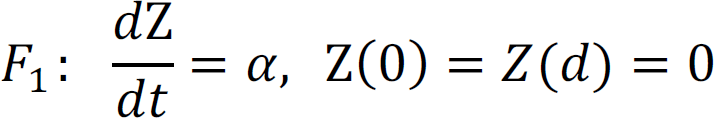

Solving this function analytically, we get the linear accumulation model:

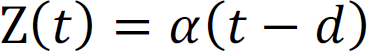

And in log-scale:

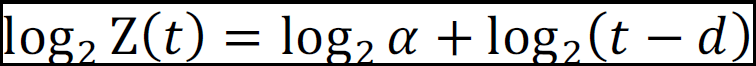

The second alternative model adds a third parameter, using a constant degradation rate (Z) to describe decay of zygotic mRNAs:

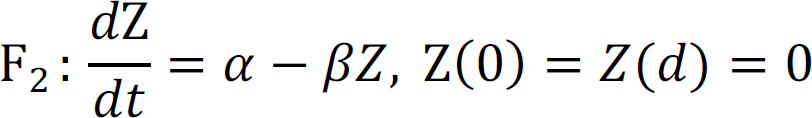

Solving this function analytically, we get:

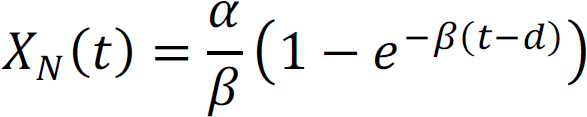

And in log-scale:

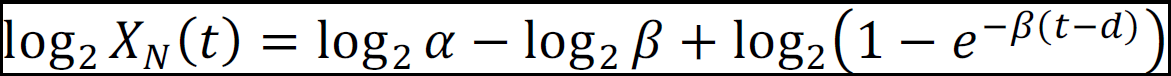

We fit each model separately to time-course data of either maternal or zygotic mRNA expression per gene. Fitting to R_STUU_ models was defined by M(t) = −4 and Q(K) = −4, and any gene that had more than 4 pseudo-bins with expression lower than -3 was also fitted to this model. Fitting to non-linear models was done using non-linear least squares regression with multiple start values with R’s *nls.multstart* package (version 1.0.0.), using 500 different start combinations (*iter=500*), and parameter bounds as listed:

|  | Start_lower | Start_upper | lower | Upper |
| --- | --- | --- | --- | --- |
| **Maternal $\beta$** | Half-life = 2 hrs | Half-life = 15 min. | Half-life = 10 hrs | Half-life = 1 min. |
| **Maternal $x_0$** | Min. expression | Max. expression | -4 | 30 |
| **Maternal $d$** | First time point with expression | Last time point with expression | First time point with expression | Inf |
| **Zygotic $\alpha$** | -15 | 1 | -20 | $10 - \log_2 60$ |
| **Zygotic $\beta$** | Half-life = 2 hrs | Half-life = 15 min. | Half-life = 10 hrs | Half-life = 1 min. |
| **Zygotic $d$** | 140 min. | First time point with expression + 60 min. | 120 min. | Inf |

Finally, we compared the fits to each of the three nested models (R_STUU_ is nested in R_Y_ which is nested in R_O_) by a likelihood ration test for each gene, and identified genes that fit R_Y_and R_O_.

The n for the likelihood ratio test was defined by:

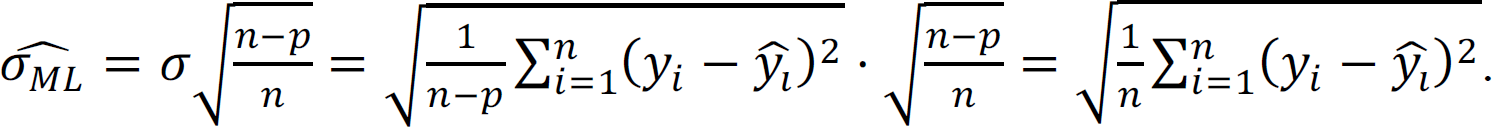

Analysis of model accuracy by goodness-of-fit test

We estimated the fit of each gene to the kinetic model using a goodness-of-fit test, by calculating:

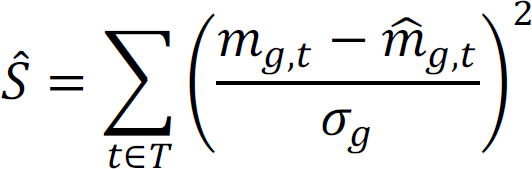

where 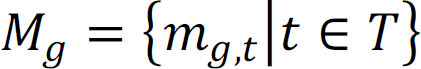 are the observed temporal samples, 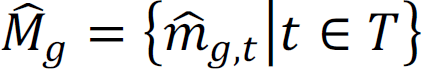 are the predictions by the model, and 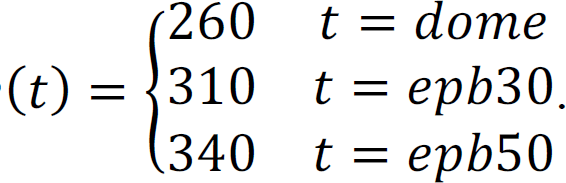

We calculated a standard deviation (n_5_) for each group of genes with a similar mean expression (ä) by comparing their expression in two replicate samples:

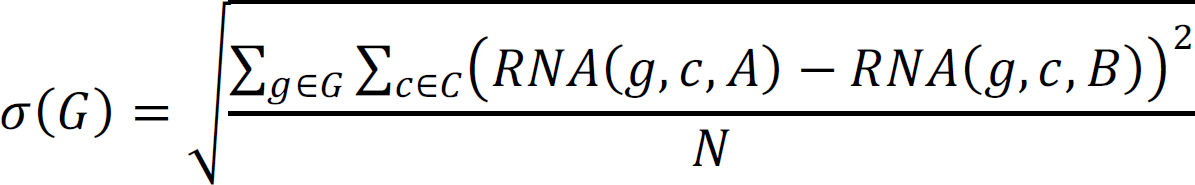

where *C* = {*epb*30, *epb*50} and *RNA(g, c, A)* is the RNA for gene g, at stage c, in each of two replicates (A or B). Genes were divided based on the quantiles of the mean expression of maternal or zygotic RNA into 6 or 10 groups, respectively. Each group of zygotic RNA contained approximately 483 genes, and each group of maternal RNA contained approximately 446 genes. We performed a chi square test 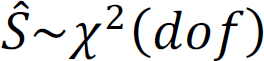, where degrees of freedom (*dof*) is the number of pseudotime bins with a reliable estimated expression for the gene.

### Analysis of model fitting by simulation studies

Simulated gene expression data was generated by selecting 5 values for each parameter in the model, resulting in 125 sets of parameters, and applying the model to generate expression data for each set, which we term “simulated genes”. For each simulated gene, we added random normal noise with mean θ = 0 and variance 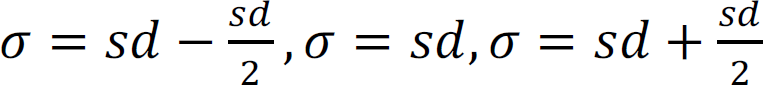, where *sd* is as described above by mean expression, and repeated the procedure 100 times for each simulated gene. The data was then fitted to the kinetic models as described above. We used a goodness-of-fit test as described above to compare the fit to the original simulated data points (without noise).

### Models of maternal and zygotic mRNA expression dynamics in developmental trajectories

For model fitting, we modified the original trajectories that were calculated during single-cell analysis. First, we combined trajectories with a more than 50% overlap in cells into a single trajectory. This way the endoderm A, B and lateral plate mesoderm were combined to a single trajectory, and the neural and nonneural ectoderm to another trajectory. Second, we excluded from the analysis trajectories that did not have estimated bulk expression for at least 6 pseudotime bins. This resulted in a final set of 5 trajectories for further analysis: enveloping layer, notochord, prechordal plate, ectoderm and mesendoderm. For each of the 5 trajectories, we calculated pseudo-bulk dynamic expression separately for cells within the trajectory and cells not assigned to this trajectory. For each of the two groups of cells, we divided the cells into pseudotime bins as described above, and applied pseudo-bulk calculation for each bin to estimate expression.

To this data, we applied two separate model fitting approaches. First, a “cell-type specific” model that assumes regulation is different within the trajectory, and thus fits two separate sets of kinetic parameters for each gene: one to fit the “trajectory-specific” profile, and another to fit the ‘non-trajectory’ profile of all other cells not assigned to that trajectory. Second, a ‘uniform’ null model, which assumes a similar rate across all cells and thus fits a single kinetic model for both the ‘trajectory-specific’ and the ‘non-trajectory’ profiles. We used a likelihood ratio test to compare the two models based on the different number of parameters used for either model, and standard variation (n) that was calculated as previously described. We calculated an effect-size for the difference between the trajectory and non-trajectory maternal-RNA and zygotic-RNA expression of each gene, by the average fold-change between the trajectory and not-trajectory expression at the three pseudo-bulk samples with maximal difference. Genes were defined as “trajectory-specific”, if the corrected p-value was smaller than 0.01, the fit to the trajectory specific model had an R-squared value above 0.8, and the effect-size was larger than 2 or smaller than 0.5.

### Grouping genes by combinations of maternal and zygotic kinetic parameters

For genes that are both maternally provided and zygotically expressed during the maternal-to-zygotic transition, we defined 4 groups of genes with distinct combinations of maternal and zygotic kinetic parameters by setting a lower-bound and upper-bound on each of the two parameters: degradation of maternal RNA (half-life bounds: < 20 pseudo-min or > 45 pseudo-min) and accumulation of zygotic RNA (log2 accumulation bounds: < -12 or = -11).

### Gene set enrichment analysis

Functional enrichment analysis was performed using *g:Profiler* R client (version e106_eg53_p16_65fcd97) with a 5% g:SCS multiple testing correction method ^44^.

Enrichment of polyA lengths ^33^ was analyzed by performing a one-sided Kolmogorov-Smirnov test (*ks.test*) between the polyA lengths of genes within a gene-set and all other genes in the dataset, using a 1% false discovery rate correction (FDR).

### Sequence k-mer enrichment analysis

3’UTR sequences were downloaded from ensembl biomart, version 103 ^45^. For each gene, annotated 3’UTRs were filtered to keep a single longest annotated 3’UTR sequence. Genes with an annotated 3’UTR sequence below 10 nucleotides were removed from the analysis. Sequences were represented by a set of all short sequences between 5-8 nucleotides long (k-mers) using *ape* and *k-mer* R packages. We associated a k-mer with a regulatory effect when genes with this k-mer in their 3’UTR had a significantly different distribution (one-sided Kolmogorov-Smirnov test, 1% FDR) of a specific parameter than genes without this sequence. We assigned an effect size to each k-mer by calculating the standardized mean difference defined as 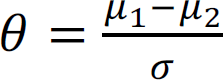, where μ_1_ is the mean μ_2_ of the first population, ô_O_ is the mean of the second population and n is the standard deviation (based on both populations).

---

**Figure S1:**
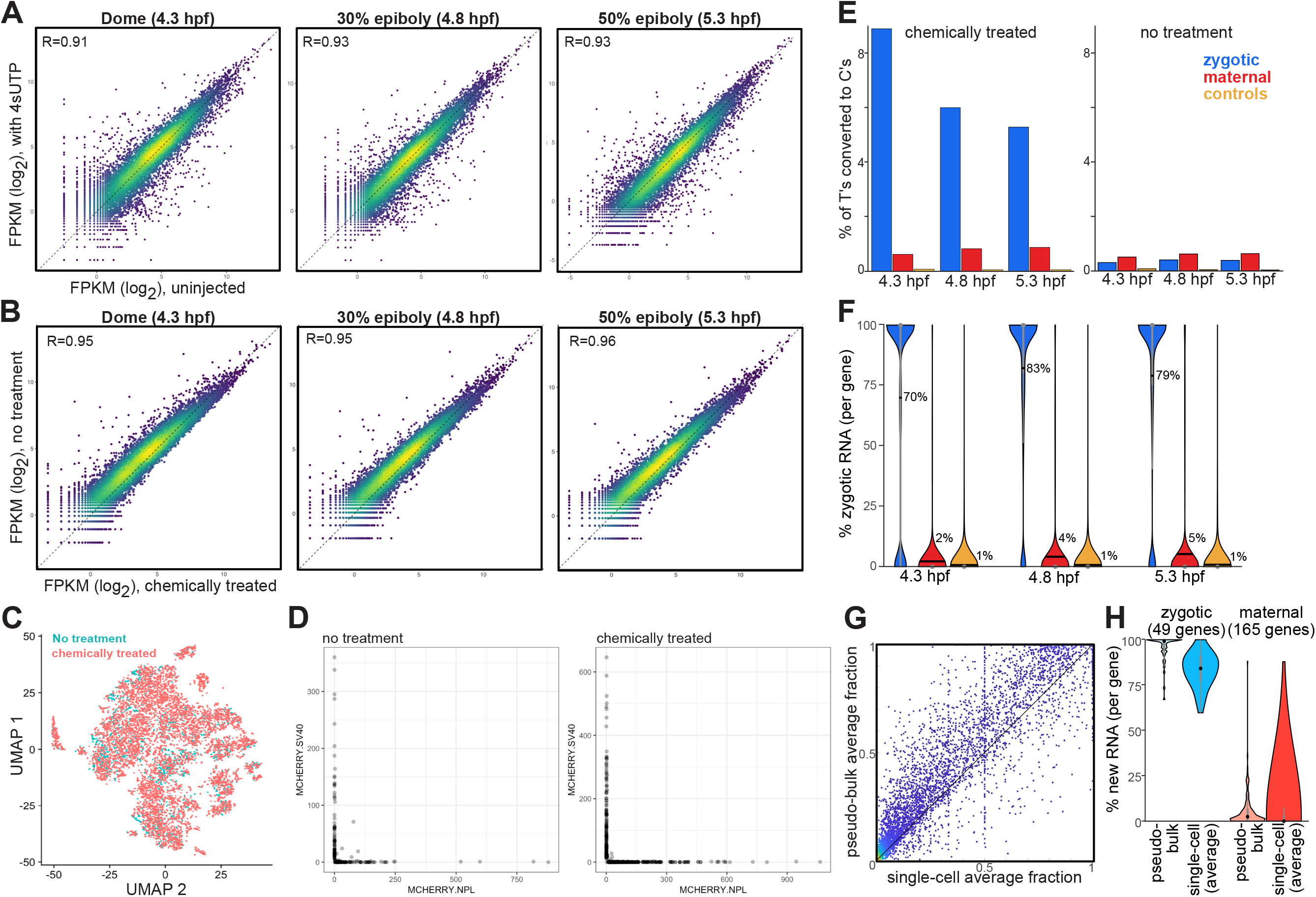
distinguishing maternal and zygotic mRNAs by combining scRNA-Seq, mRNA metabolic labeling and nucleotide conversion in zebrafish embryos. **(A-B)** Scatter plots of RNA expression (log_2_) in 3 embryonic developmental stages (left to right: dome, 4.3 hpf; 30% epiboly, 4.8 hpf; 50% epiboly, 5.3 hpf). Colors represent density (yellow=high density; blue=low density). Pearson R values are indicated on plots. Expression was calculated as the normalized sum of all UMIs in cells within each stage. **(A)** scRNA-Seq results of uninjected embryos (x-axis; data from ^11^) or embryos that were injected with 4sU-triphosphate (4sUTP) at the 1-cell stage (y-axis; this study). **(B)** scRNA-Seq results of embryos that were injected with 4sU-triphosphate (4sUTP) at the 1-cell stage, and were either chemically treated (x-axis) or not treated (y-axis). **(C)** UMAP projection of single cells from embryonic samples of embryos that were injected with 4sU-triphosphate (4sUTP) at the 1-cell stage, and were either chemically treated (red) or not treated (cyan). **(D)** Scatter plots of UMI counts associated with each single-cell in 3 out of 5 of our samples that included a mixture of embryos that were randomly injected with one of two mCherry mRNA species (bearing a different sequence at its 3’UTR), labeled as MCHERRY.NPL (x-axis) and MCHERRY.SV40 (y-axis). Samples were either chemically treated (right) or not treated (left). Less than 0.05% of cells in both converted and non-converted samples were associated with reads from both mCherry species, indicating that conversion did not interfere with RNA attachment to beads and did not increase barcode mixing. **(E)** Fraction of T bases that were sequenced as C (y-axis), within each of 3 temporal samples (x-axis), when applying chemical treatment (left) or without such treatment (right), calculated for subsets of 110 known zygotic transcripts (blue), 332 known maternal transcripts (red), and 3 injected control mRNAs without any incorporated 4sU residues (yellow). As label was introduced in the 1-cell stage, transcripts from zygotic genes (blue) are expected to be fully labeled in all samples. Differences in converted fraction can arise from different incorporation efficiencies between samples, different position-specific incorporation rates, genetic polymorphism and other confounding effects. **(F)** GRAND-SLAM estimates of per-gene percent of newly-transcribed zygotic RNA (y-axis) within each single cell, for subsets of 100 known zygotic transcripts (blue), 288 known maternal transcripts (red), and 3 injected control mRNAs without any incorporated 4sU residues (yellow). Some of the genes analyzed in panel (E) were not analyzed by GRAND-SLAM due to relatively low expression within single cells. The central dot is the median; black line is the average (value is also noted on plot); the edges of the gray box are the 25th and 75th percentiles. GRAND-SLAM analysis reduced biases in converted fractions, resulting in similarly estimated fractions of labeled and unlabeled RNA across samples. **(G)** Scatter plot of the fraction of zygotic mRNA assigned to genes by either average of single cell estimations (x-axis) or a pseudo-bulk calculation (y-axis). Colors represent density (yellow=high density; blue=low density). **(H)** Violin plots of the distribution of the fraction of zygotic mRNA assigned to 49 known zygotic genes (left, blue) and 165 known maternal genes (right, red) by either average of single cell GRAND-SLAM estimations (dark colored) or a pseudo-bulk GRAND-SLAM calculation (light colored). The central dot is the median; the edges of the gray box are the 25th and 75th percentiles. The characteristically low label incorporation rates in combination with low per-cell number of reads by scRNA-Seq, limited the accuracy of estimated labeled mRNA fraction within single cells. The estimated labeled mRNA fraction for known zygotic genes was often lower than expected within single cells, resulting in an unlabeled mRNA background. By aggregating single cells by stage and applying GRAND-SLAM estimation to these pseudo-bulk samples we increased the accuracy of estimated labeled mRNA fraction, bringing the estimated fraction for known zygotic genes to nearly 100%.

**Figure S2:**
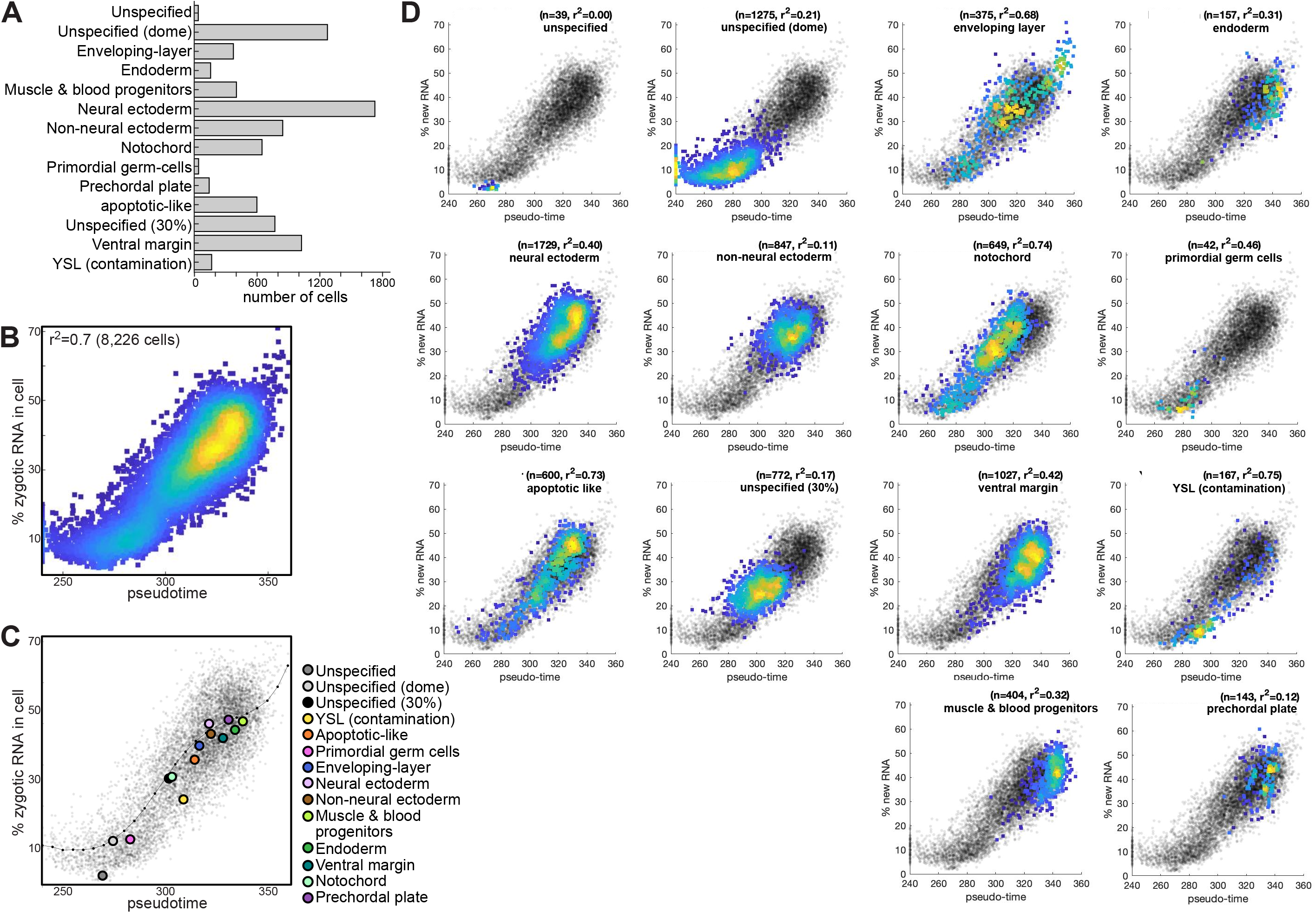
fraction of newly-transcribed zygotic mRNA in single cells is correlated to pseudotime. **(A)** Histogram of number of cells (x-axis) assigned to each of 12 distinct cell-types that were defined in the data (y-axis). Endoderm A + B were merged. **(B-D)** Scatter plots of pseudotime assigned to cells (x-axis, pseudo-min) and zygotic mRNA fraction in cells (y-axis). A cell specific fraction of zygotic RNA (across all genes) was calculated by the ratio of zygotic UMI counts to total UMI counts in the cell. **(B)** Colors represent density (yellow=high density; blue=low density) across all cells in dataset. Pearson R-squared value is indicated on plots. **(C)** Colored dots represent the averages of all cells in each cell-type, and are colored by cell-types (as indicated). Black dots and line represent a pseudotime sliding window average of fraction of zygotic mRNA in cells. **(D)** Black dots represent all cells in dataset. Colors in each plot represent density (yellow=high density; blue=low density) for cells from a specific cell-type, as indicated. Pearson R-squared value and number of genes are indicated on each plot.

**Figure S3:**
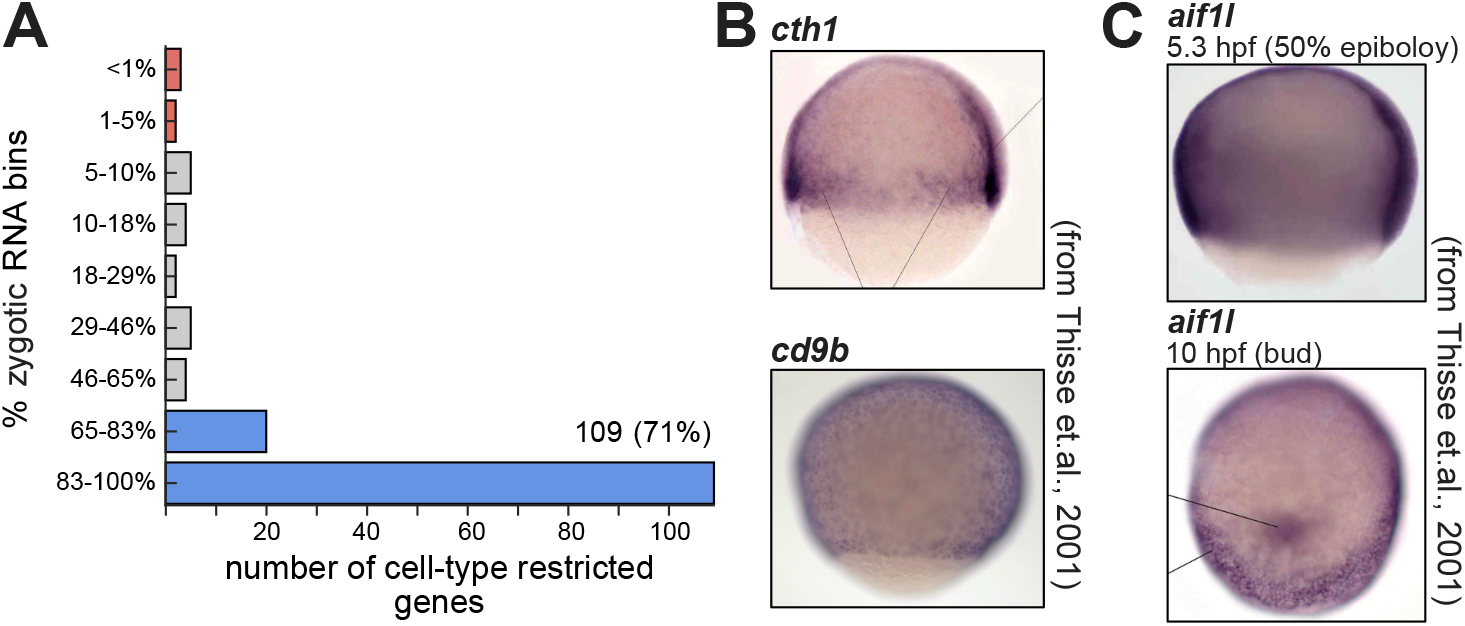
variability in zygotic mRNA accumulation between genes. **(A)** Histogram of genes with cell-type restricted expression (y-axis, number of genes) by their % of zygotic RNA quantile (9 quantiles, x-axis). Bars corresponding to top two bins are considered zygotic genes (blue, zygotic RNA fraction >65%) while bottom three bins are considered maternal genes (red, zygotic RNA fraction <5%). **(B-C)** In-situ hybridization staining of zebrafish embryos for mRNAs of different embryonic genes, at specific developmental stages, as indicated. Images from ^46^.

**Figure S4:**
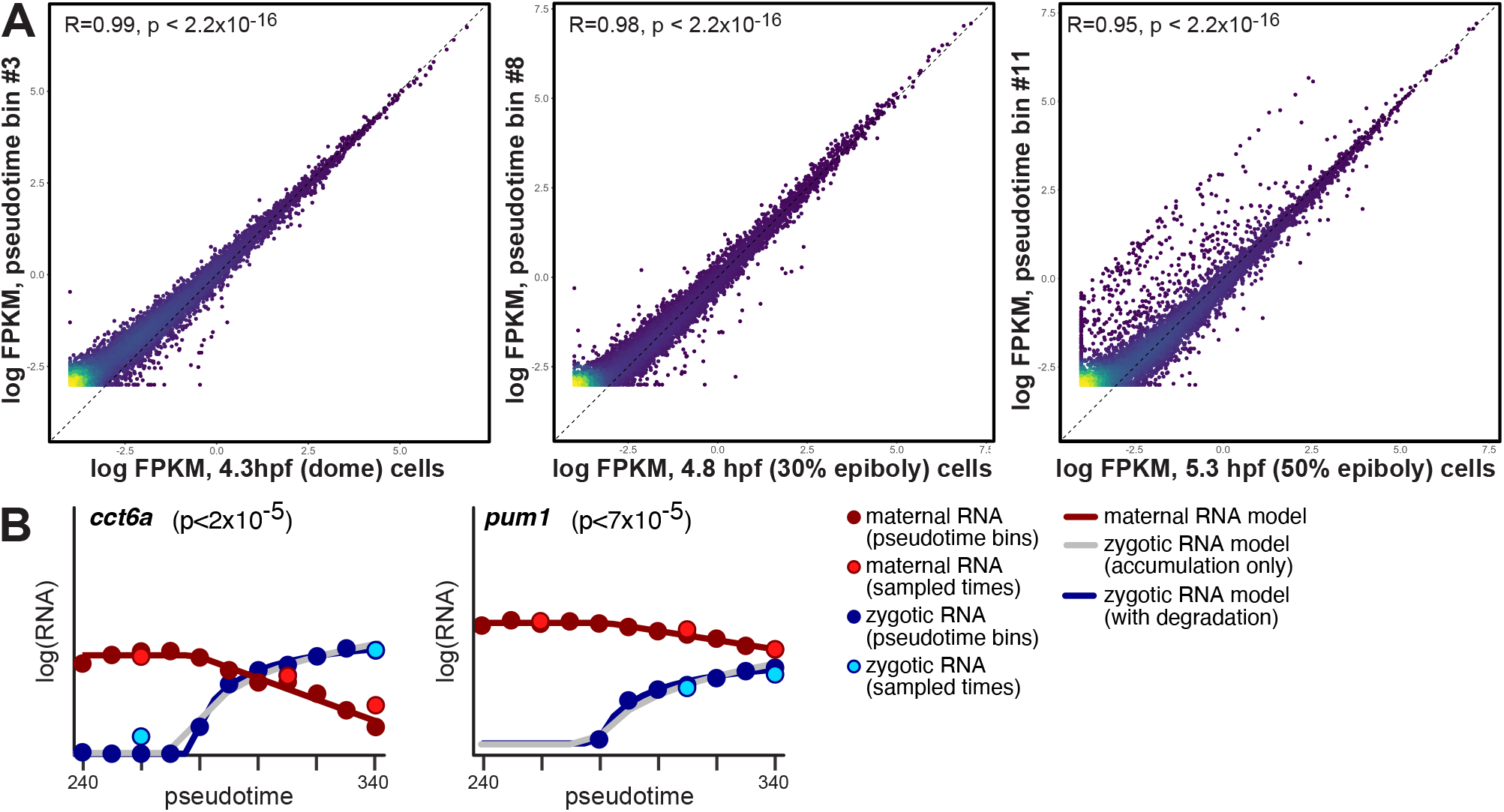
modeling the dynamics of the maternal and zygotic transcriptomes. **(A)** Scatter plots of RNA expression (log_2_) 3 embryonic developmental stages (x-axis; left: dome, 4.3 hpf; middle: 30% epiboly, 4.8 hpf; right: 50% epiboly, 5.3 hpf) and in 3 pseudotime bins (y-axis; left: bin #3; middle: bin #8; right: bin #11). Colors represent density (yellow=high density; blue=low density). Pearson R values are indicated on plots. **(B)** Model fits (solid lines) to interpolated zygotic (blue dots) and maternal (red dots) expression levels (y-axis, log_2_ scale) across 11 pseudotime (x-axis) bins for genes which significantly rejected the zygotic accumulation model (gray line) in favor of the more complex model with zygotic degradation (blue line). Gene name and likelihood ratio test p-values (for selecting between simpler or complex model) are indicated on top. Light blue and light red dots represent estimated zygotic and maternal mRNA levels, respectively, for three sampled timepoints (4.3 hpf, 4.8 hpf and 5.3 hpf).

**Figure S5:**
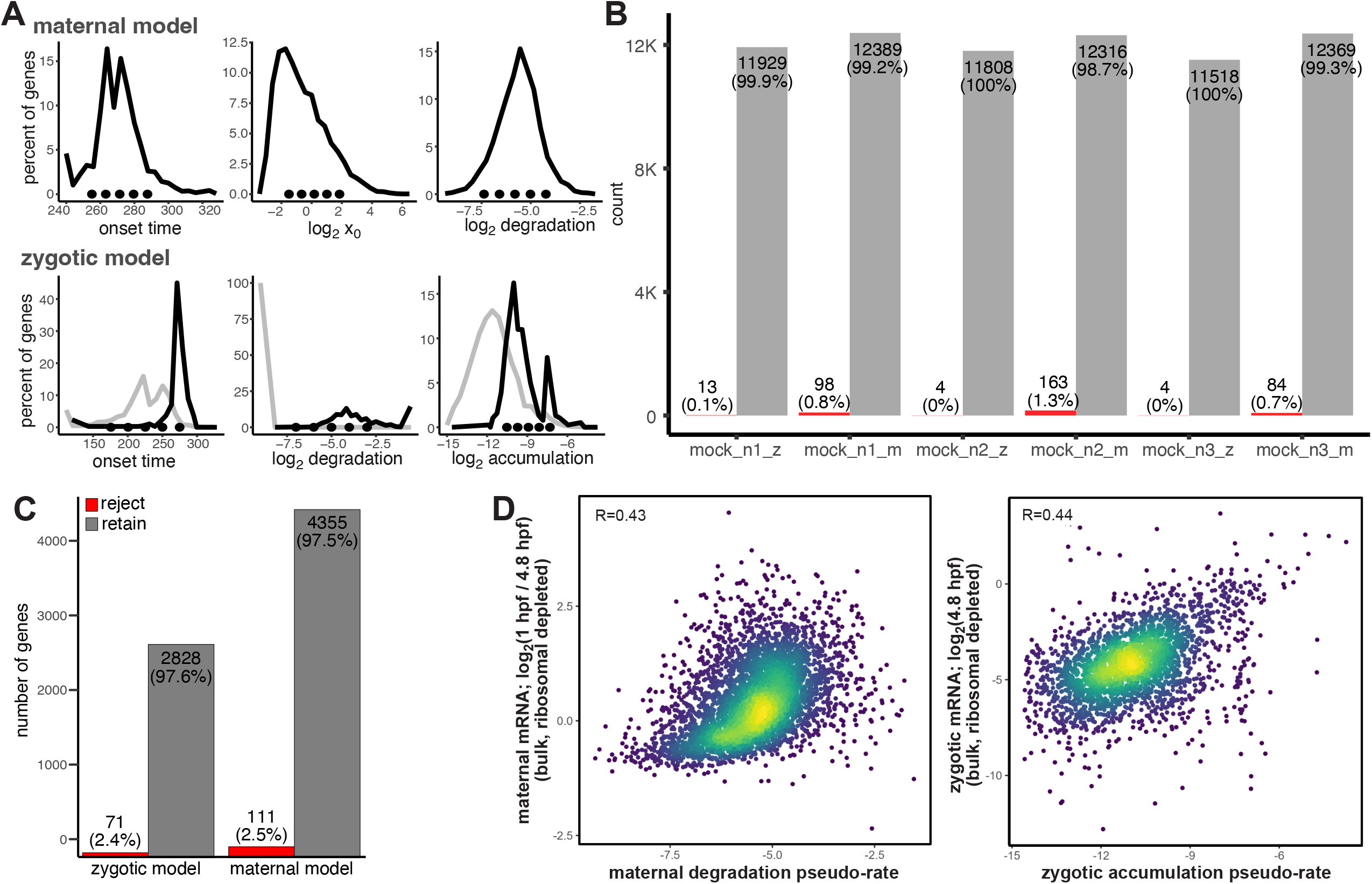
validations of kinetic models for maternal and zygotic mRNA expression dynamics. **(A)** Selection of parameter values for simulation studies for each of 6 model parameters. Shown are distributions of parameter values that were fitted by the maternal or zygotic RNA expression models to genes. Top: maternal decay model parameters (y-axis, density; x-axis left to right: degradation onset time (pseudo-min), initial expression level (log_2_), degradation pseudo-rate (log_2_ 1/pseudo-min)). Bottom: zygotic accumulation model parameters, either without a degradation term (black), or with a degradation term (gray) (y-axis, density; x-axis left to right: transcription onset time (pseudo-min), degradation pseudo-rate (log_2_), accumulation pseudo-rate (log_2_ RNA/pseudo-min)). **(B)** Histogram of number of mock generated samples (y-axis) that retained (gray) or rejected (red) the fitted model by a goodness-of-fit test in each of 3 simulation experiments (x-axis) with increasing noise levels: 50% of standard deviation (as estimated in replicated samples) (n1), 100% of standard deviation (n2) and 150% of standard deviation (n3). Number of samples is indicated. **(C)** Histogram of number of genes (y-axis) that retained (gray) or rejected (red) the fitted model (x-axis) by a goodness-of-fit test (chi-square p-value). Number of genes is indicated. **(D)** Scatter plots comparing kinetic parameters that were estimated for each gene by the model (x-axis; left: maternal mRNA degradation pseudo-rate, right: zygotic mRNA accumulation pseudo-rate) to fold-change of mRNA that was estimated in ribosomal depleted bulk samples (y-axis, log_2_; left: maternal mRNA, right: zygotic mRNA). Fold change of maternal mRNA was estimated by ratio of expression in 1 hpf (4-cell) to 4.8 hpf (30% epiboly) samples. Fold change of zygotic mRNA was estimated by the expression in 4.8 hpf (30% epiboly) sample, since zygotic mRNA expression at the 1 hpf sample was below detection. Colors represent density (yellow=high density; blue=low density). Pearson R values are indicated on plots.

**Figure S6:**
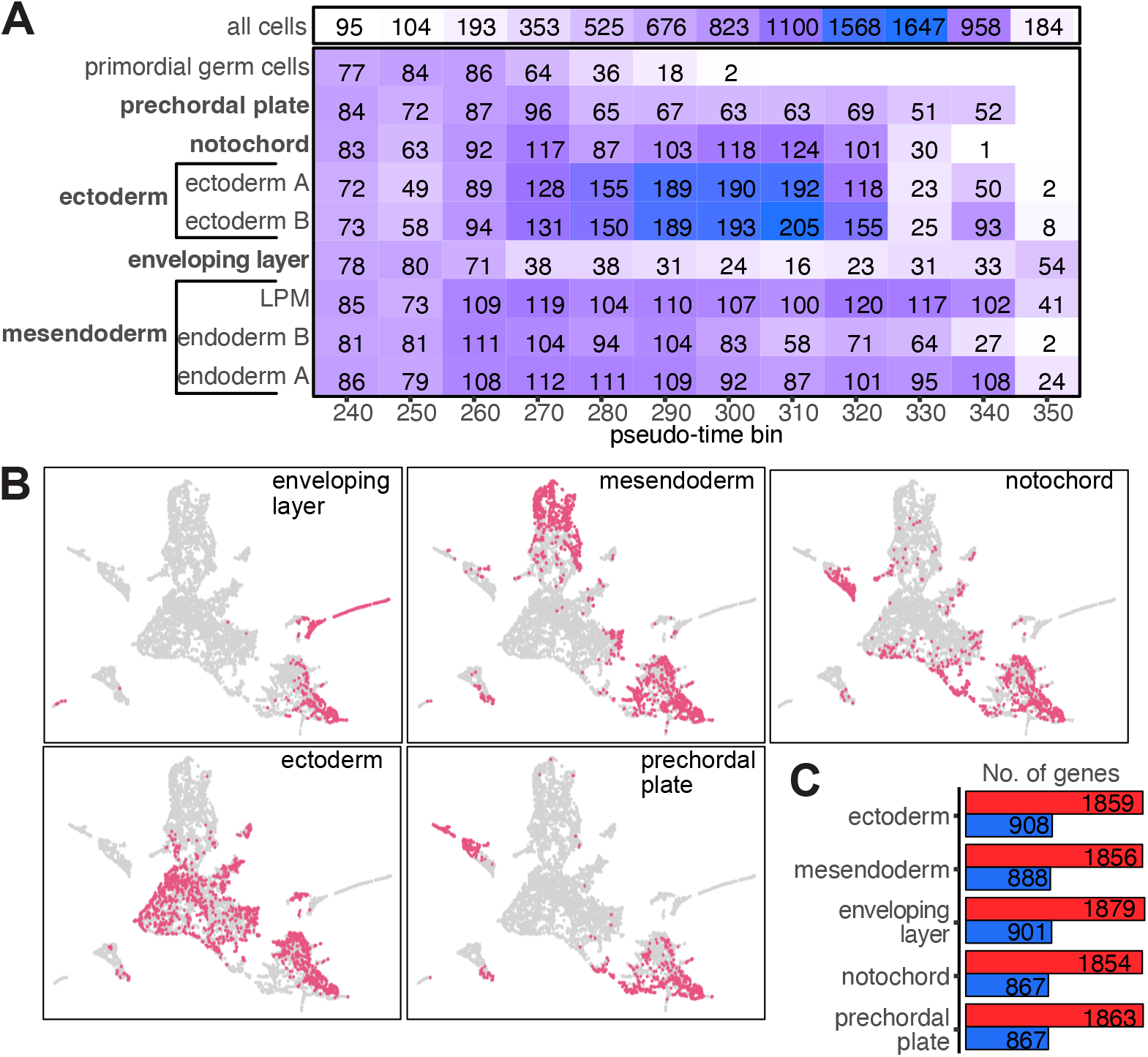
modeling mRNA regulation within developmental trajectories. **(A)** Number of cells in each subset of cells (rows) assigned to 11 pseudotime bins (columns). Color-scale represents number of cells (white = no cells; dark purple = maximal number of cells). Number of cells is indicated. **(B)** Single-cell UMAP projection with cells assigned to each of 5 developmental trajectories (as indicated on map) colored in red. **(C)** Histogram of number of genes (y-axis) analyzed in each developmental trajectory (x-axis). Red: maternal genes, Blue: zygotic genes.

**Figure S7:**
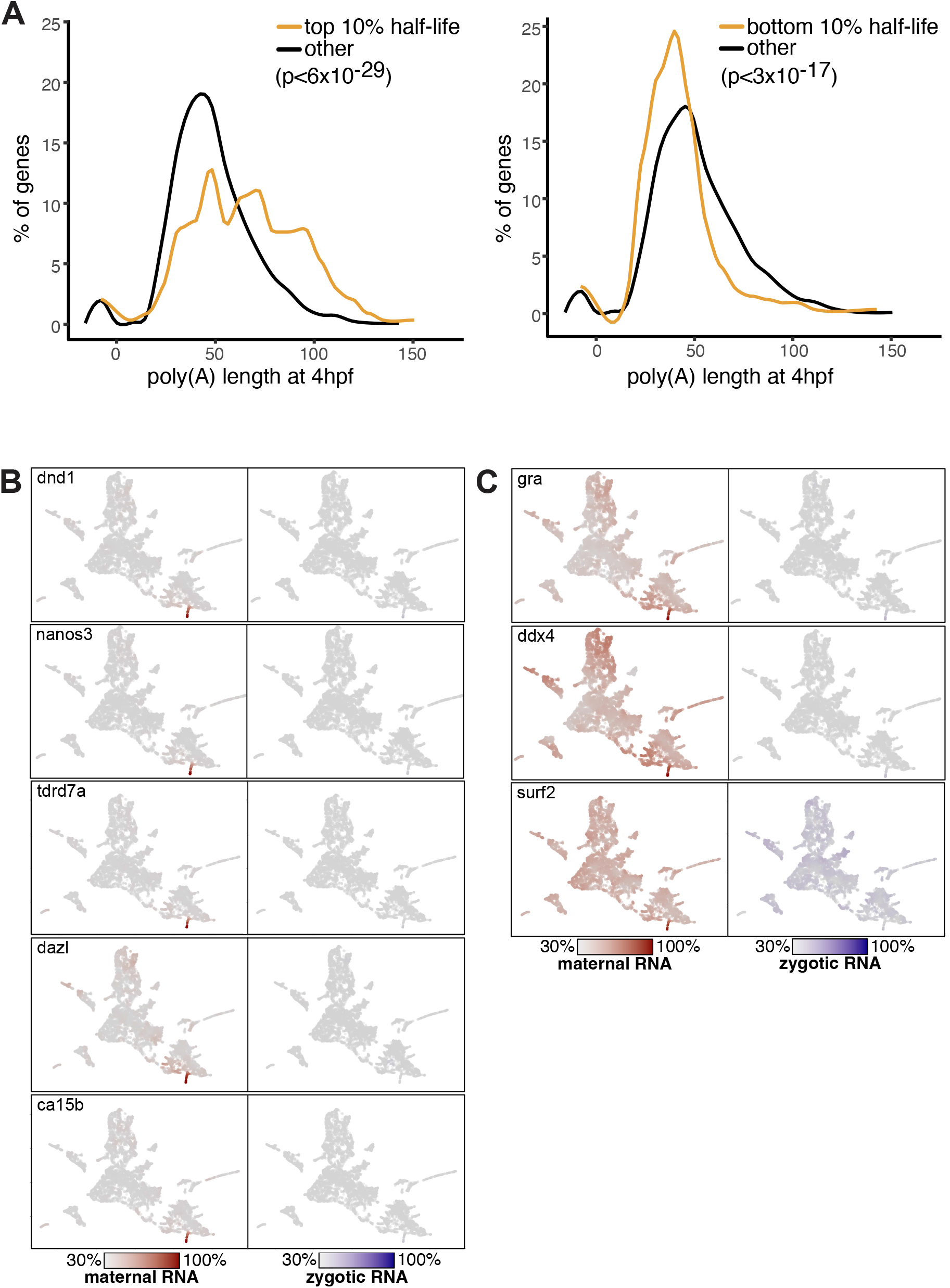
sequence enrichments associated with regulatory rates of mRNA degradation. **(A)** Frequency (y-axis) of poly(A) tail length (x-axis) as measured by ^33^. Left: for genes with half-life at the top 10% (yellow) or other (black). Right: for genes with half-life at the bottom 10% (yellow) or other (black). P-value of a one-sided Kolmogorov-Smirnov test (FDR <1%) is indicated. **(B-C)** Single cell expression of 8 germ-cell specific expressed genes across all 8,226 collected single cells. All cells are plotted, and each cell is colored by the normalized expression of a gene’s maternal copies (red, left) or zygotic copies (blue, right). Color-scale of each gene is scaled by its maximal total expression, and its minimal 30% quantile of maternal and zygotic RNA expression. Analyzed genes are indicated on plot. Plotted genes show cell lineage specificity within germ-cells with either **(B)** low somatic background or **(C)** high somatic background.

**Table S1:**
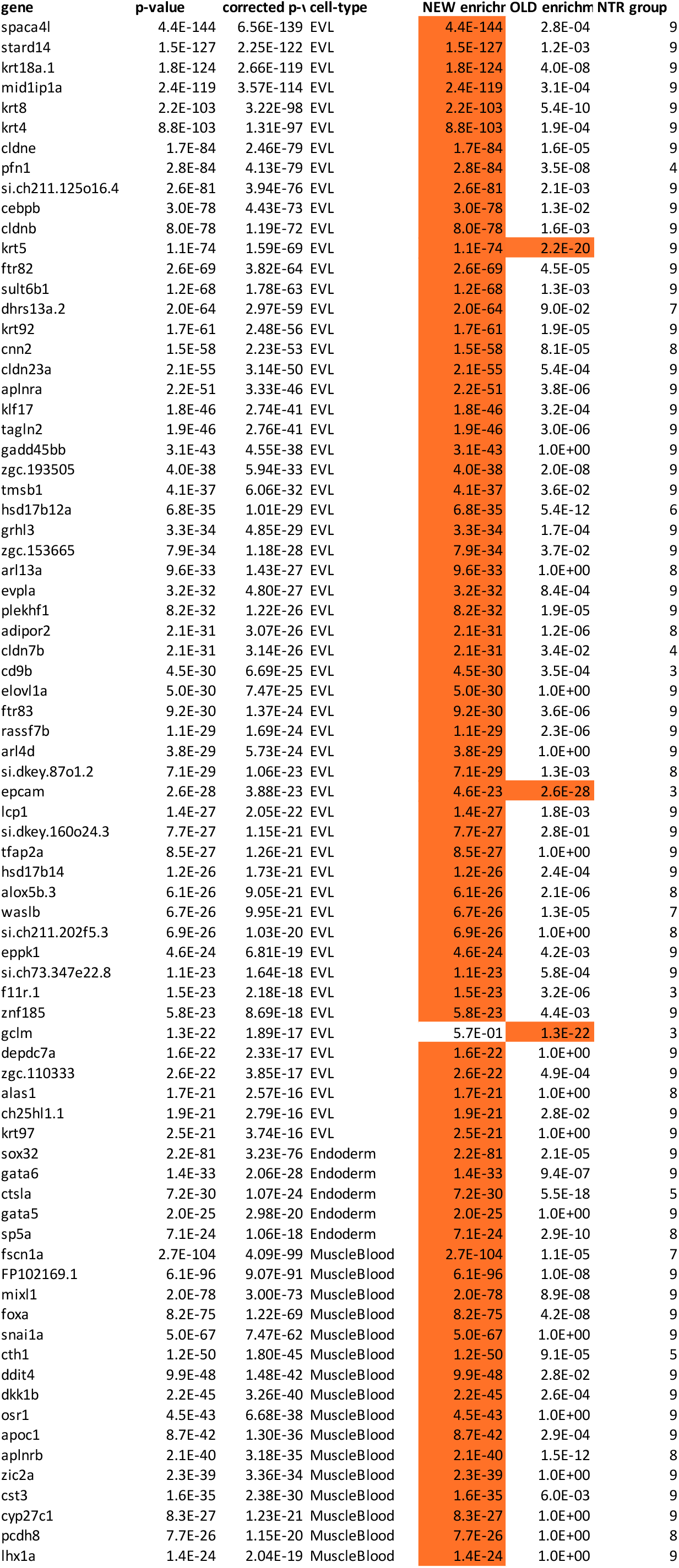
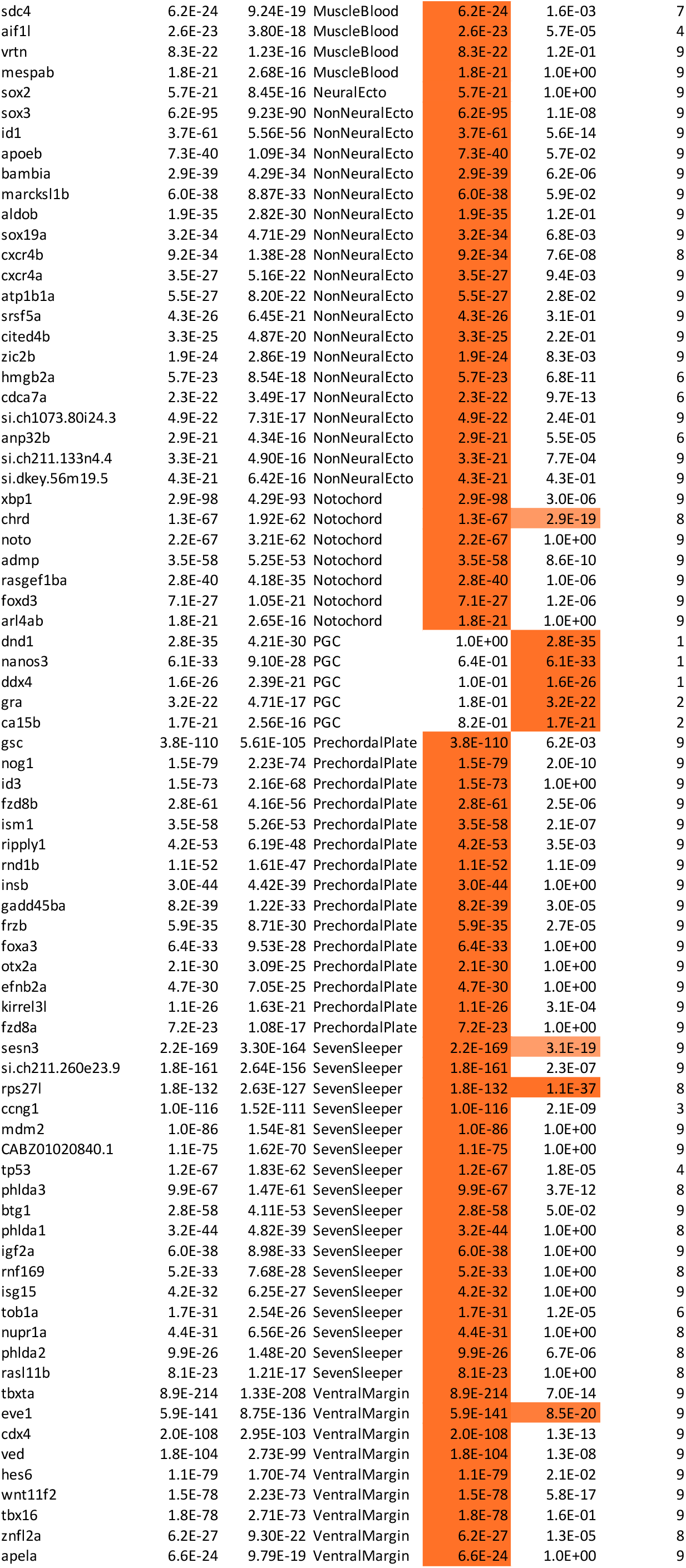
List of 436 cell-type enriched genes.

**Table S2:**
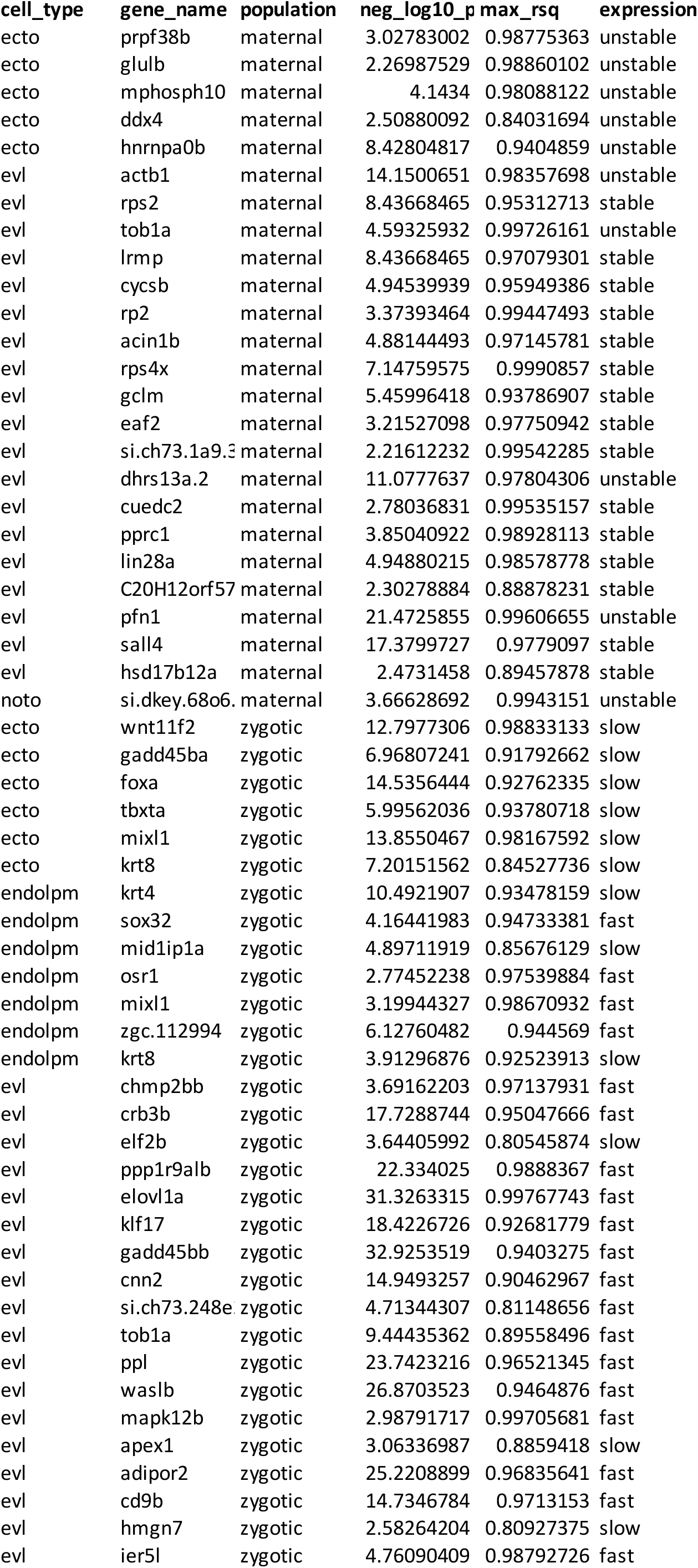
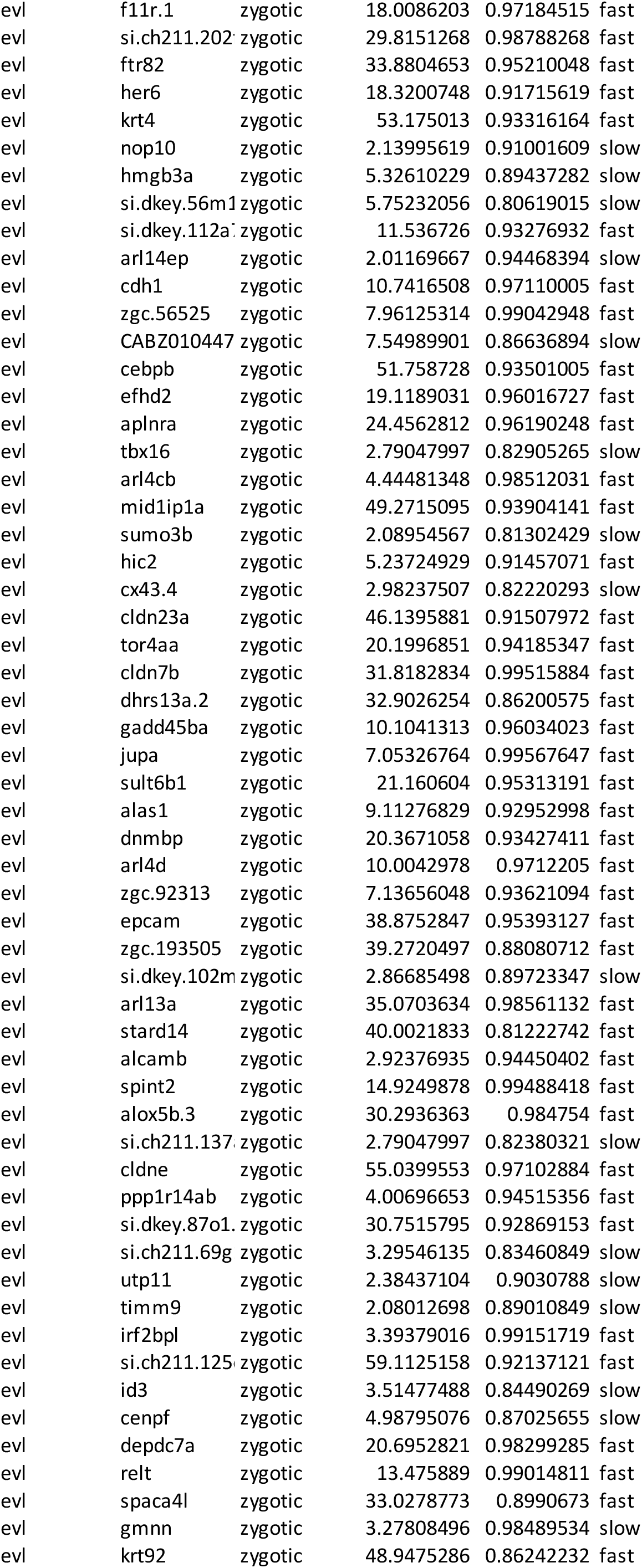
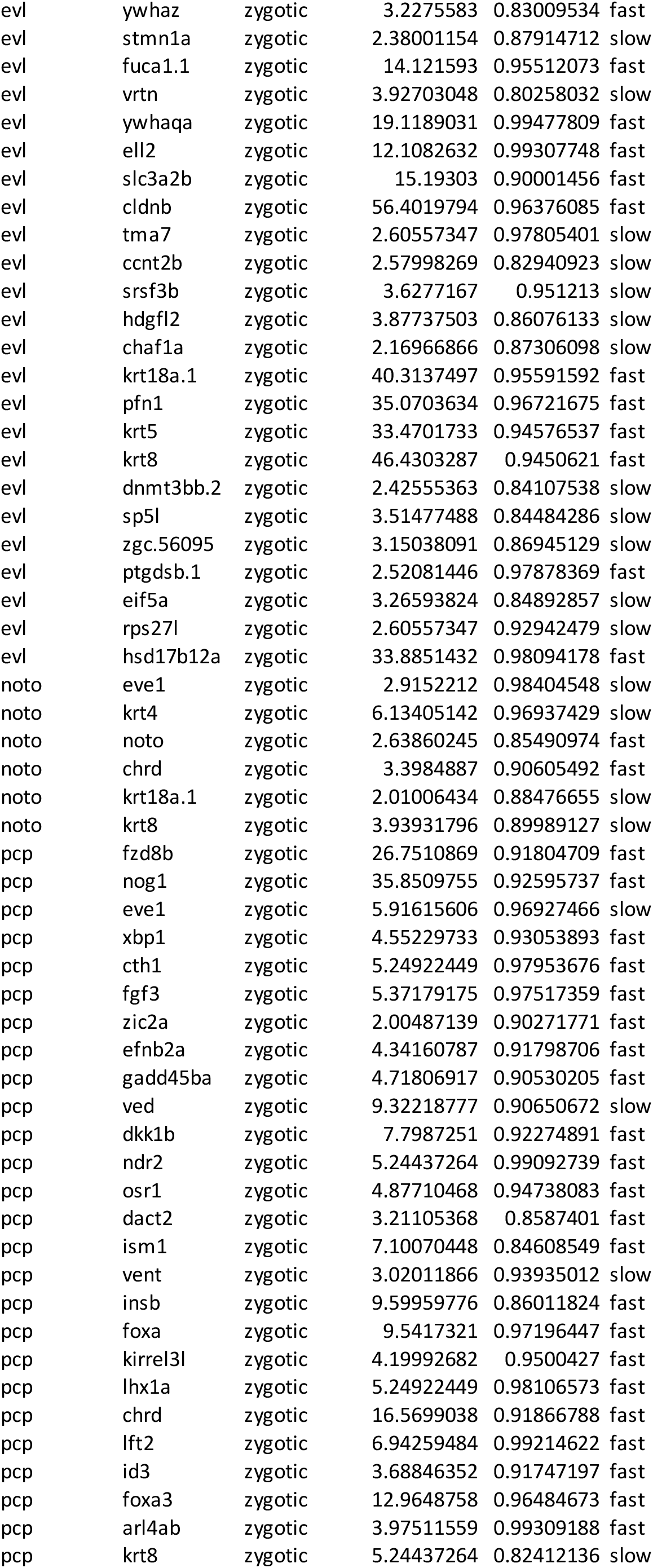
List of 149 genes with trajectory-specific mRNA dynamics.

